# Complementary killing activities of *Pbunavirus LS1* and *Bruynoghevirus LUZ24* phages on planktonic and sessile *Pseudomonas aeruginosa* PAO1 derivatives

**DOI:** 10.1101/2025.04.09.647956

**Authors:** Maud Billaud, Clarisse Plantady, Benoît Lerouge, Emma Ollivier, Julien Lossouarn, Elisabeth Moncaut, Julien Deschamps, Romain Briandet, Aurore Cleret, Cindy Fevre, Gaëlle Demarre, Marie-Agnès Petit

## Abstract

Four *P. aeruginosa* phages active against a representative panel of strains, and with complementary spectra of action were chosen with the goal of using them for phage therapy. Two of them were myoviruses belonging to the *Pbunavirus LS1* species, and two were podoviruses belonging to the *Bruynoghevirus LUZ24* species. In order to better apprehend the interactions of these phages with their *P. aeruginosa* host bacteria, we undertook the characterization of their bacterial receptors, using a PAO1 derivative as a recipient strain. Whereas the receptor of the *P. LS1* phage Ab27 had already been characterized as the O-antigen chain of the lipopolysaccharides, no information was available at the onset of this work on the receptor used by the phages of the *B. LUZ24* species. We show that the surface polysaccharide Psl is this receptor. Psl stands for <u>p</u>olysaccharide <u>s</u>ynthesis locus, and it is an important component of the biofilm matrix in a large panel of *P. aeruginosa* strains, including PAO1. Remarkably, the *B. LUZ24* phages were more active against PAO1 in minimal medium compared to rich medium. Consistently, this was correlated with larger amounts of Psl bound at the bacterial surface during exponential growth in the minimal medium compared to the rich medium. Biofilms formed on a medical intubation device, as well as in in 96-well plates, were degraded to different extent by the two phage species: biofilms grown for 7 hours on tubing device were degraded more efficiently by the *B. LUZ24* than the *P. LS1* phage, whereas mature biofilms (16 hours) formed in 96-well plates were degraded more rapidly by *P. LS1* than by *B. LUZ24* phage. The frequency of genetic mutants resisting to each phage were determined in liquid medium by a fluctuation assay and found in the range of 10^-5^ to 10^-6^ per generation. Interestingly, most of the *P. LS1* resisting mutants were more sensitive to the *B. LUZ24* phage. We conclude that the combination of the four selected phages has very promising properties, which should be relevant in the framework of phage therapy.

## INTRODUCTION

*Pseudomonas aeruginosa* is a ubiquitous Gram-negative bacterium found in freshwater and terrestrial environments. It is also an opportunistic pathogen in humans, causing bloodstream, urinary tract, or burn wounds infections, as well as airways infections in cystic fibrosis patients or patients under mechanical ventilation (Jonckheere et al., 2018; Ramirez-Estrada et al., 2016). Among its many virulence factors, the three polysaccharides alginate, Psl, and Pel are particularly important, as they permit the establishment of chronic biofilm infections and immune evasion (Jones and Wozniak, 2017), which is the main cause of ventilator associated pneumonia (VAP). Some clinical isolates (such as PAO1) will build up their biofilm mostly on Psl, whereas others (such as PAK1) mostly on Pel, and a third category on a mix of both Pel and Psl (Colvin et al., 2011; Jones and Wozniak, 2017). In all cases however, Psl seems needed for the attachment step of the biofilm, and monoclonal antibodies targeting Psl have been tested for adjunctive use to antibiotics, to treat biofilms (Ray et al., 2017). Psl polysaccharide is a neutral polymer composed of repeating pentasaccharides of D-mannose, D-glucose, and L-rhamnose, with a cell associated- and a cell-free form (Byrd et al., 2009). How the Psl polysaccharide is attached to the bacterial surface remains unknown at present. Regulation of its synthesis is multi-layered and relies in particular on the intracellular levels of di-cyclic-GMP (Dreifus et al., 2022; Feng et al., 2020; Zhu et al., 2016). Interestingly, even during planktonic growth in liquid medium, Psl-dependent aggregates tend to form in early exponential growth (OD 0.05-0.2) and to dissociate later on (Melaugh et al., 2023).

Besides its effectiveness as a pathogen, *P. aeruginosa* belongs to the ESKAPE group of pathogens for which multi-drug resistance is becoming a threat. Indeed, *P. aeruginosa* infections are getting more difficult to treat, since fewer antibacterial drugs are available, resulting in higher mortality and morbidity rate (Boucher et al., 2009). There is therefore an urgent need for novel therapeutic strategies, such as phage therapy (Forde and Hill, 2018; Pires et al., 2020).

Similar to the outcome of antibiotic treatments, bacterial populations placed under phage attack lead to the selection of clones resisting to phage. This has been observed under *in vitro* settings, but also in the mouse gut (Cornuault et al., 2020; De Sordi et al., 2017), as well as in some cases of patients treated with phage therapy over long periods of time (Schooley et al., 2017). In short-term *in vitro* experiments with *P. aeruginosa*, bacterial resistance to phage was found to consist either in acquiring CRISPR spacers against the phage, or mutating the phage receptor (Alseth et al., 2019; Hesse et al., 2020; Yang et al., 2020).

To maintain phage infectivity regardless of bacterial evolution, using different phages (both in terms of taxonomy and bacterial receptor) is a good strategy (Kunisch et al., 2024). It should slow down the arising of phage resistant bacteria in the patient submitted to phage therapy. One can expect that the generation of bacterial mutants modifying two different receptors is less likely to happen (Yang et al., 2020). Similarly, strains (or populations of bacteria) acquiring spacers against different phages are less likely to emerge. Moreover, it was demonstrated that the CRISPR-Cas system is associated with an infection-induced fitness cost: although mutants acquiring spacers are initially selected for, the receptor mutants are selected in the long-term (Meaden et al., 2021).

For *P. aeruginosa*, most characterized phages are using a limited set of receptors, such as the lipopolysaccharide (LPS) or the type IV pili (Ceyssens and Lavigne, 2010; Harvey et al., 2018; Yang et al., 2020). In addition, phage OMKO1 (φKZ-like) was found to use another receptor, OprM, an outer membrane porin. Interestingly, this porin is also used by two multidrug efflux pumps, MexAB and MexXY (Chan et al., 2016). In fact, some of these receptors can also be considered virulence, or fitness factors. When bacterial resistance emerge, resistant clones are often counter-selected, or have fitness trade-offs, making it harder for them to survive under *in vivo* conditions (Forti et al., 2018; Kortright et al., 2019). A case of phage resistant mutations affecting efflux pumps was also found to lead to the resensitization of the strain to antibiotics (Chan et al., 2016). The use of phages targeting virulence factors as receptors could make phage-therapy doubly effective: in addition to killing a large part of the bacterial population, it may drive multi-drug resistant bacterial pathogens to evolve towards phage resistance at the cost of decreasing their fitness and ability to survive in the host. Thus, it is important to identify the bacterial receptors of virulent phages that might be used in phage therapy.

A large screening of *P. aeruginosa* phages active against an international panel of *P. aeruginosa* strains (De Soyza et al., 2013) had led us to select four phages with complementary spectra of action, with the goal of using them for phage therapy. Two of them were myoviruses belonging to the species *Pbunavirus LS1* (below, these two phages will be collectively referred to as *LS1* phages), and two were podoviruses belonging to the species *Bruynoghevirus LUZ24* (*LUZ24* phages below). Whereas the primary receptor of the *LS1* phage Ab27 had already been characterized as being the O-antigen part of the LPS (Latino et al., 2016), no information was available at the onset of this work on the receptor used by *LUZ24* phages (Ceyssens et al., 2008). We therefore undertook this characterization, which led us to uncover a new receptor for *P. aeruginosa* phages, the Psl polysaccharide. We also found that both phage species were able to degrade biofilms, with different kinetics and final efficiency depending on the biofilm support.

## MATERIALS AND METHODS

### Bacterial Strains, Plasmids and Culture Conditions

Bacterial strains and plasmids used and constructed in this study are listed in. Table S1. Unless indicated, *E. coli* and *P. aeruginosa* strains were grown at 37°C, 200 rpm in Luria Bertani Broth containing 5 g/L NaCl (Formedium), which will be designated below as Lennox. The minimal medium used was ‘Modified MOPS Medium’ with 20 mM acetate, designated below MMMa (LaBauve and Wargo, 2012). Compared to the usual MOPS medium (Neidhardt et al., 1974), CaCl2 and FeCl2 were added. Sputum Medium was prepared as described in (Palmer et al., 2007). To grow strains in partial aerobiosis, the Lennox medium was covered with 2 mL of sterile paraffin over the 10 mL culture, and incubated without agitation in a 50 mL sterile plastic tube. Moreover, KNO3 was added at 100 mM to favor *P. aeruginosa* anaerobic growth (Pallett et al., 2019).

For *E. coli* strains, gentamicin (Gm) was added when required at 30 µg/mL. For *P. aeruginosa* strains, antibiotic and chemicals were added at the following concentration: isopropyl β-D-1-thiogalactopyranoside (IPTG): 0.5 mM, and Gm: 10 µg/mL for pSV35 derivatives, and 50µg/mL for pSEVA629M containing strains.

Bacterial strains were stored at −80°C in Lennox or MMMa, with 20% glycerol.

### Construction of strain PAO1-3Δ

Strain PAO1_OR (accession LN871187) contains two prophages, Pf4 and Pf6. Unless otherwise stated, a PAO1_OR derivative in which these prophages were inactivated was used throughout this work and named PAO1-3Δ. Starting from PAO1_OR, deletion of Pf4, of the *rep* gene of Pf6, and of *toxA*, were performed successively by allelic exchange, using plasmids pMB01, pMB03 and pMB06, respectively (Table S1). Each deletion involved three steps (unless specified otherwise). In the first step, a non-replicative pEX18ApGW derivative in which the region to be deleted was replaced by an *aac1* gene, and flanked by ∼300 bp regions homologous to the bacterial chromosome was introduced by single crossing-over into the chromosome (with a gentamicin resistance selection). In a second step, the deletion of the plasmid while retaining the deletion was selected by plating bacteria on sucrose and gentamicin, as the presence of the *sacB* gene on the pEX18ApGW plasmid renders the strain sensitive to sucrose. In a third step, the *aac1* gene, which is flanked by FRT sequences, was excised with plasmid pFlp2. For the *toxA* gene deletion, to avoid introducing a third FRT scar, no *aac1* gene was placed between the regions flanking the deletion, and vector pEXG2 was used instead of pEX18ApGW. The construction involved only the two first steps, and *toxA* mutants were screened by PCR. The final strain PAO1-3Δ was completely sequenced and assembled. The three constructed deletions were as expected. Two additional mutations were generated during the construction steps, a 1 bp deletion within the gene *PAO1OR3357* encoding an hypothetical protein, and a CAC deletion within a run of 15 CAC repeats of the *czcD_2* gene, coding for a cadmium cobalt and zinc antiporter.

### Plasmid constructions

Primers used in this study are listed in Table S2. All PCR were done using Phusion High Fidelity DNA Polymerase (New England BioLabs). All plasmid cloning steps were performed in *E. coli* strain JM105. To generate the fragments needed for the cloning of pMB01, pMB03 and pMB06, each of the two or three PCR fragments to be assembled were first amplified separately, and then a splicing overlap extension PCR allowed to join the fragments together. Final plasmid construction integrity was verified by Sanger sequencing.

To complement *pslA* mutants, the *pslA* ORF and its 20 preceding nucleotides including the RBS was PCR amplified on DNA from PAO1-3Δ, using primers pslA_5’ and pslA_3’ (Table S2), and the PCR product was digested by *Kpn*I and *Eco*RI and cloned into the *Kpn*I/*Eco*RI digested plasmid pPSV35, downstream of the Plac promoter, to generate pPSV35-*pslA*. Integrity of the cloned ORF was verified by Sanger sequencing. The same approach was used to clone the *pslD* ORF in pPSV35 (primers pslD_5’ and 3’), and generated plasmid pPSV35-*pslD* (Table S2).

Most plasmids were introduced into PAO1 strains by conjugation. For this, the plasmid was first transformed into strain β2163, which contains in its chromosome all genes needed for the RP4 conjugation process. Its growth depends on the addition of 0.3 mM of diaminopurine. Next, the plasmids, containing the RP4 oriT transfer origin, were transferred into the recipient strain by conjugation, as described in (Vallet-Gely and Boccard, 2013). The pGEX2 derivative, as well as pSEVA629M plasmid was introduced into *P. aeruginosa* strains by electroporation.

### Phages

The four phages used in this study, PP1450 (species *P. LS1*), PP1777 (species *P. LS1*), PP1792 (species *B. LUZ24*) and PP1797 (species *B. LUZ24*) were isolated from wastewater samples. Their genomes were sequenced and functionally annotated. For this, coding sequences were first predicted with RAST (Aziz et al., 2008) using genetic code 11, virus option and RASTtk pipeline. The predicted proteins were then annotated in parallel with different tools, and then annotated by consensus. Tools included (i) PsiBLAST search against the Conserved Domain Database (Lu et al., 2020) with an E-value cutoff of 10^-5^, (ii) remote homology search of structural genes against the Virfam database (Lopes et al., 2014), (iii) global remote homology search using HHblits against the PHROGs database (Terzian et al., 2021), and (iv) remote homology search with the HHPred interface against the PDB, Pfam-A_v35 and UniProt-SwissProt_viral70_3_3_Nov_2021 databases (Zimmermann et al., 2018). In all cases, biological function predictions were retained if HHsearch probability was above 95%. The genome alignments of Figure 1 were generated with BLASTn and drawn using GenoplotR (Guy et al., 2010).

**Figure 1:**
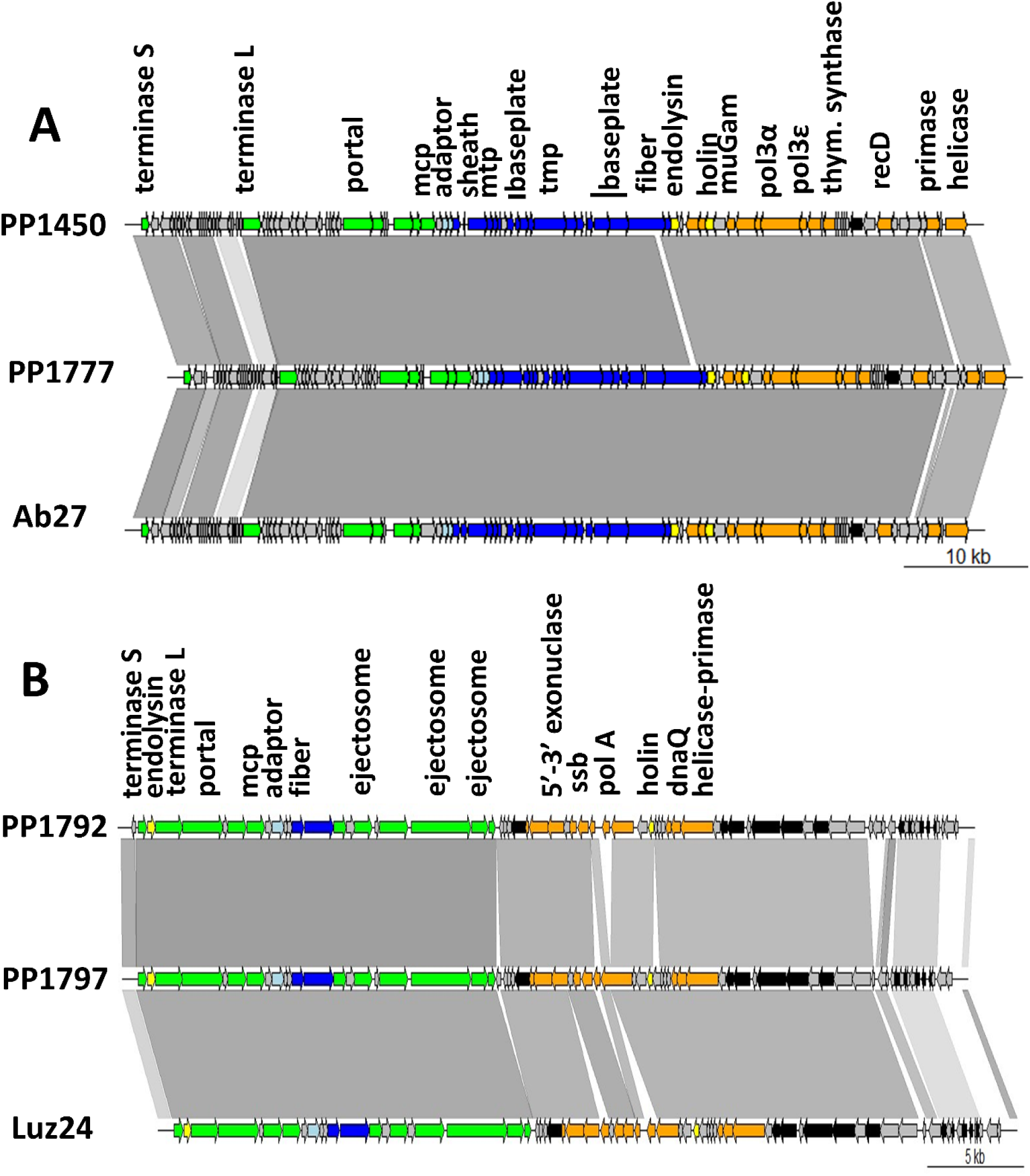
**A.** Genomic alignment of phages PP1450 and PP1777 with Ab27 (orientation and first nt of PP1450 and PP1777 were adapted, to align with Ab27). **B**. Genomic alignment of phages PP1792 and PP1797 with Luz24 (orientation and first nt of PP1792 and PP1797 were adapted, to align with Luz24). Genes are colored according to their functional modules. Green, capsid and encapsidation; light blue, connector; dark blue, tail; orange, DNA replication; black, auxiliary metabolic genes, grey, hypothetical proteins. The gradient of grey between the phage maps represents the BLASTn identity value (90% to 100% nt identity range).

### Growth monitoring of bacteria during phage infection

For large volume cultures, overnight cultures were diluted to OD_600_=0.05 in fresh medium. At OD_600_ =0.5, corresponding to ∼10^8^ colony forming unit (CFU)/mL, 5 mL of bacterial cultures were infected with phage at a multiplicity of infection (MOI) of 1, incubated with no agitation for 10 minutes at 37°C, then diluted 10-fold in 50 mL of pre-warmed Lennox medium, and incubated at 37°C for several hours with shaking, in a 250 mL flask. The OD_600_ was followed over time. Culture growths in 96-well plaques were monitored as follows: overnight cultures were diluted to ∼10^6^ CFU/mL in fresh medium. Phages were prepared at ∼10^10^ PFU/mL and serially diluted. Bacteria and phage solutions were added to each well in a final volume of 150 µL, exposing bacteria (∼10^5^ CFU/mL) to different phages concentrations (∼10^1^ to 10^9^ PFU/mL). The last two columns served as positive controls (bacteria without phages) and blank (only media). Plates were incubated at 37°C while shaking. OD600nm was measured every hour for 24 hours using a MultiSkan Go or a LogPhase 600 plate reader.

### Fluctuation assay with the Luria Delbrück test (Luria and Delbruck, 1943)

Fluctuation assays were performed either in Lennox or MMMa. Overnight cultures were diluted 100-fold and incubated at 37°C with shaking. At OD_600_ =0.5, the culture was diluted to 10^6^ CFU/mL, dispatched into 90 wells of the 96 well plate (100 µL) and mixed with 10^8^ PFU (100 µL) of phages. The six remaining wells contained: the growth medium (2 wells); the growth medium with phage and no bacteria (2 wells); the growth medium with tested bacterial strain and no phage. The plate and its plastic cover were then wrapped in parafilm to avoid evaporation and incubated at 37°C, 200 rpm for 24 h for Lennox growth, and 36 h for MMMa growth. After this incubation time, the number K of clear cultures, i.e. in which all bacteria were killed (meaning that in this well, no resisting mutant was present at the time of infection) was monitored. If K was between 20 and 80, we considered that the fraction K/90 was an estimation of the first term of the Poisson law with mean λ (K/90 = e^-λ^), describing the occurrence of a mutation with low frequency in the population. Mutation frequency *f* was then obtained by dividing λ by the average number of bacteria per well N, at infection time (*f* = - Ln (K/90) / N).

### Selection of bacterial strain resistant to phage, sequencing, and mutation analysis

Bacterial mutants were recovered either at the end of the small volume fluctuation assays, or after overnight growth of large batch cultures of phage infection experiments. For large batch cultures, the culture the next morning was serially diluted 10^6^ and 10^7^-fold and plated. For fluctuation test assays, three wells with bacterial growth per experiment were chosen randomly and streaked on agar plates containing the same growth medium. Each mutant was streaked three times on either Lennox or MMMa to eliminate residual phages. After the purification process, mutants were grown overnight in Lennox or MMMa at 37°C, 200 rpm and stored at - 80°C with 20% glycerol.

DNA from mutants resistant to *LS1* or *LUZ24* phages was extracted, and either entirely sequenced (Illumina Hiseq) or sequenced only for the genes of interest by Sanger technology. Shotgun sequencing was performed on an Illumina Hiseq platform (Eurofins Genomics, 2 x 125 bp, depth: 9 million reads). Mutations were identified on reads using Breseq (Deatherage and Barrick, 2014), with PAO1-3Δ as the reference genome.

### Spot test and efficiency of plating assay (EOP)

The plating efficiency of the four phages on *P. aeruginosa* mutant strains was determined by spot assay: 10 μL of a serial 100-fold dilutions of a phage preparation were spotted on double-layer agar plates inoculated with each bacterial host. The number of plaques observed after overnight incubation was compared to the phage titre on its reference strain: PAO1-3Δ for PP1450 and PP1777; NAR71 for PP1792 and PP1797.

### Adsorption assay

Bacteriophage adsorption assay was performed as previously described (Kropinski, 2009). Briefly, overnight bacterial cultures incubated at 37°C were diluted in fresh medium at OD_600_ of 0.05 and incubated at 37°C until reaching OD_600_= 0.5. Phages were then added at MOI 0.01, and the adsorption proceeded at 37°C for 10 minutes without shaking. Then, samples were collected and either centrifuged at 10 000 *g* for 5 minutes or treated with 50 µL of chloroform (both methods gave comparable results), and supernatants were titrated by spot test. As a reference, a negative control, with phage only and no bacteria, was performed and titrated.

### Surface Psl extraction and Psl immunoblots

Surface Psl extraction was performed following the protocol described in (Byrd et al., 2009). Briefly, for all samples an equal number of bacteria, equivalent to 5 mL of a culture at OD 2, was centrifuged 10 min at 10 000 *g*. For cultures grown in rich medium, to remove residual Psl from supernatants, bacteria were washed twice with fresh Lennox, by centrifugation. The clean pellet was then resuspended in 200 µL EDTA 0.5 M, boiled at 95°C for 20 min. The bacterial debris were then removed by centrifugation 5 min at 16000 *g*, and the supernatant was treated with proteinase K at 0.2 mg/mL and incubated 1h at 60°C. Finally, proteinase K was inactivated by incubation of the samples at 80°C for 30 min. Psl extracts were kept at 4°C, for a maximum of 1 week before testing. No treatment was applied to recover Psl from culture supernatants. Bacteria were simply pelleted by centrifugation (5000 *g*, 5 min), and the supernatant was filtered with 0.2 µm filters and kept at 4°C.

To quantify Psl, 5 µL of samples were deposited on a 0.45 µm nitrocellulose membrane (Amersham Protran ref 10600003), and let dry. Membranes were then incubated in TBST buffer (50 mM Tris pH 7.5, 150 mM NaCl, 0.1% Tween 20) complemented with 1% fat-free powder milk (Regilait Bio) for 1 hour in a rotating shaker at room temperature, in a volume of 5 mL. Then human anti-Psl monoclonal antibody (Creative Biolabs. Ref MRO-160MZ) was added at a 1/5000 dilution, and incubation was continued 1 hour. After 3 short rinses of the membrane with TBST, the secondary antibody (Goat-anti-human IgG antibody coupled to HRP, Sigma, Ref AP112P) was applied in 5 mL TBST buffer, at 1/5000 dilution, and incubated for 1 hour under the same shaking and temperature conditions. After 3 short rinses of the membrane with TBST, the revealing reagent (Clarity Western ECL substrate, BioRad ref 170-5060) was poured on the membrane, which was imaged 1 minute later, in a Chemidoc apparatus (BioRad). Image analysis was performed with the ImageLab software of BioRad, using non-saturated images and volume counts adjusted with local background. Three-fold dilutions of the most concentrated Psl sample allowed to determine that Psl quantities were measurable in a range of 27-fold under these conditions.

### Biofilm quantification on medical device

The intubation device (Hexamed, reference 44295LCM/93350LCM) used to grow biofilms were prepared as follows: balloons, bevel connections and the small injection tube were removed; the remaining tubing was cut in 1 cm long cylinders. Each cylinder was then cut into 4 longitudinal sections and each piece was placed in a well of a 96 U-shaped wells plate (Thermofisher scientific, reference 167425). Wells were seeded with 150 µl of an overnight culture of the tested bacterial strain diluted 1/1000 in Lennox. Plates were covered and incubated for 7 h without shaking at 30°C. Each intubation piece was then washed by 3 successive immersions in PBS and placed in a clean 96 well plate. Phages were added at a final concentration of 10^9^ PFU/mL in 150 μL Lennox, covered and incubated for 17 h at 30°C without shaking.

Crystal violet coloration was performed as follows: each intubation piece was washed by 3 successive immersions in PBS and placed in a 96 deep well plate (VWR, reference 732-3325) containing 500 μL of 0,01% of crystal violet (CV) solution, and incubated for 30 min at RT without shaking. The excess of crystal violet was removed by 3 successive immersions in PBS to allow biofilm visualization. Cristal violet was then dissolved in 500 μL of 33% acetic acid for 10 min with shaking and quantified by OD at 590 nm. After subtraction of the blank (33% acetic acid), the test OD was normalized by the intubation piece weight. Each experiment included triplicates, and a point on the graph represents the average of the 3 phage-treated replicates divided by the average of the untreated condition.

### Quantification of biofilm growth in 96-well plates by confocal microscopy

Strain MB198, a PAO1_OR derivative in which plasmid pSEVA627M (Silva-Rocha et al., 2013) was introduced, was always propagated in Lennox complemented with 50 µg/mL gentamicin. Its constitutive GFP expression permitted to follow living bacteria within the biofilm. To detect dead bacteria, propidium iodide (PI, Invitrogen) was used (16-80 µM final concentration). Prior microscopy analysis, a growing or a mature biofilm was grown during 6 and 16 hours, respectively, at 37°C, under static condition, in a 96-well plate (microscopic-grade Phenoplate-96, Perkin Elmer). To inoculate each well, an overnight culture of MB198 was diluted to OD 0.05 in Lennox, and a 200 µL volume was applied to the well, and incubated 1 hour at 37°C statically, to let bacteria adhere. Next, the 200 µL were removed and replaced by fresh Lennox, and placed back in the incubator. After biofilm growth, phages were added at 5×10^6^ and 5×10^8^ PFU per well, for the short and long pre-incubations respectively, in a 50 µL volume of SM buffer (50 mM Tris pH 7.5, 100 mM NaCl, 10 mM MgSO4) mixed with PI; un-infected wells received 50 µL of SM buffer mixed with PI, and the plate was placed under the microscope. The phage addition constituted time 0 of the experiments. Confocal imaging was performed using a HCS-SP8 Leica Confocal laser scanning microscope (LEICA Microsystems, Germany) at the INRAE MIMA2 microscopic platform (doi.org/10.15454/1.5572348210007727E12). A 3D time-lapse acquisition was initiated, capturing image stacks every hour for a total duration of 48 hours. Each image stack was acquired with a 1µm z-step interval. Images (512×512 pixels covering a 184.52 µmx184.52 µm area) were captured at a 600 Hz frequency using a 63x water objective lens with a numerical aperture of 1.2. For GFP detection, excitation was set at 488 nm and emission was collected in the 500-550nm range. For PI, excitation was also at 488 nm, with red fluorescence captured in the 600-750 nm range. 2D projections were generated using Imaris 9.3.1 software (Bitplane, Zurich, Switzerland). Quantitative biofilm parameters, including biovolume, thickness and roughness, were extracted using BiofilmQ software (Hartmann et al., 2021).

## RESULTS

### The four virulent phages infecting *P. aeruginosa* selected for phage therapy belong to two phage species, Pebunavirus LS1 and Bruynoghevirus LUZ24

Four *P. aeruginosa* infecting phages were selected for therapeutic use, based on their efficient lytic capacities and good coverage of an international reference panel of *P. aeruginosa* strains (De Soyza et al., 2013). Their genomes were sequenced and compared to phage genomes at NCBI (BLASTn). Two of them, PP1450 (accession LV539990) and PP1777 (LZ998055) were closely related to the 66 kb Ab27 genome (Latino et al., 2016), a myovirus belonging to the *Pbunavirus LS1* species. They shared with Ab27 98.11% and 97.41% nucleotide identity, respectively, over 97% of their genome length (Figure 1A). We concluded they belonged to the *LS1* species, like Ab27. The other two phages, PP1792 (LZ998056) and PP1797 (LZ998057) were closely related to the 45 kb Luz24 genome (Ceyssens et al., 2008), a podovirus of the *Bruynoghevirus LUZ24* species. They shared with Luz24 97.02% and 96,59% nucleotide identity, respectively, over 94% of their genome length (Figure 1B). They belonged therefore to the *LUZ24* species.

Two phages per species were chosen for therapeutic use, because despite their partial overlap, they had interesting complementary host ranges. We therefore searched for variable regions within putative receptor binding proteins. Among the large genomic changes visible between the two *LS1* genomes, there was a short region within a tail fiber gene that may correspond indeed to the receptor binding protein. No similar large modification was observed in the tail nor the fiber gene of the two *LUZ24* genomes (dark blue genes in Figure 1), but some amino-acid polymorphism was detected in the fiber gene (Figure S1), which may also be at the root of the differing host ranges of PP1792 and PP1797.

None of these four phages encoded an integrase gene, confirming their virulent lifestyle. No toxin, nor antibiotic resistance gene were detected in these genomes, ensuring them to be safe for therapeutic use.

### Mutants resisting to LS1 phages arise at a frequency of 6 to 7 x 10^-6^ per generation

To study phage host interactions, a PAO1_OR derivative deleted of its prophages was constructed (see Methods for strain construction) and designated PAO1-3Δ. To estimate the frequency at which mutants resisting to each *LS1* phage arise, fluctuation assays were performed. For this, PP1450 or PP1777 were propagated at high MOI on a given concentration of PAO1-3Δ bacteria, in Lennox medium in 90 replicate cultures, and the number of cultures in which mutants arose allowed to determine mutation frequency, as described in the Method section. This frequency was 6.1 (±4.1) ×10^-6^ per generation for PP1450, and 7.0 (±1.7) ×10^-6^ for PP1777.

### The receptor of the two LS1 phages is the O-antigen of the lipopolysaccharide

We next proceeded to characterize the obtained mutants. For this, wells in which bacterial growth emerged were selected, and mutants isolated by 3 successive streaks on Lennox medium. For five independent mutants (3 resisting to PP1450, 2 resisting to PP1777), complete genome sequencing was performed and reads were then analyzed to search for mutations (Table 1).

**Table 1:** Mutations detected in the genomes of PP1450- or PP1777-resistant mutants.

| Strain name | Selection conditions | Gene mutated/lost | Detected mutation | Protein modification |
| --- | --- | --- | --- | --- |
| MB79 | Fluctuation assay with PP1777 | *wzy1* | A:7→6 (nt 136) | T54* |
| MB75<sup>§</sup> | Fluctuation assay with PP1450 | *wzy2* | A:7→8 (nt 136) | R74* |
| MB81 | Fluctuation assay with PP1450 | *wzy2* | A:7→8 (nt 136) | R74* |
| MB76 | Fluctuation assay with PP1450 | *wzy3* | G:6→7 (nt 625) | T276* |
| MB78 | Fluctuation assay with PP1777 | Numerous, including *galU* | Δ 329 240 bp |  |

<sup>§</sup>Mutant MB75 had an additional mutation in the *gloA2* gene (R50H).
\*Indicates the stop codon

Four of these five mutants had a mutation in the *wzy* gene, which encodes a polymerase involved in the biosynthesis of the LPS O-antigen chain, resulting in truncated proteins (Table 1). The last mutant had a large deletion including the *galU* gene, which is also needed for the biosynthesis of LPS. Its defect results in the absence of the O-antigen chain and a defective outer core. This type of large deletion has been observed repeatedly with the PAO1 strain, including among mutants resisting to phage infection (Shen et al., 2018; Yang et al., 2020). We noted that the two *LS1* phages had different plating efficiencies on the mutants: PP1450 was more affected than PP1777 on *wzy* mutants, whereas conversely, PP1777 was more affected than PP1450 on the large deletion mutant MB78 (Table S3).

Mutant strain MB79 (Table 1), with a deletion of an A in a run of 7 A resulting in a truncated protein of 54 amino acids (mutation *wzy1*), was kept for further study. We verified that compared to PAO1-3Δ, the adsorption of PP1450 on MB79 was significantly decreased (Figure 2, p-value < 0.0005). Sequencing of the *wzy* gene PCR product in 6 additional resistant clones also revealed mutations in this gene, so that in total among 11 sequenced mutants, 10 were in the *wzy* gene, and one was a deletion including *galU*. Altogether these results suggest that, in agreement with results obtained for Ab27 (Latino et al., 2016), the O-antigen chain LPS is the receptor of phages PP1450 and PP1777.

**Figure 2:**
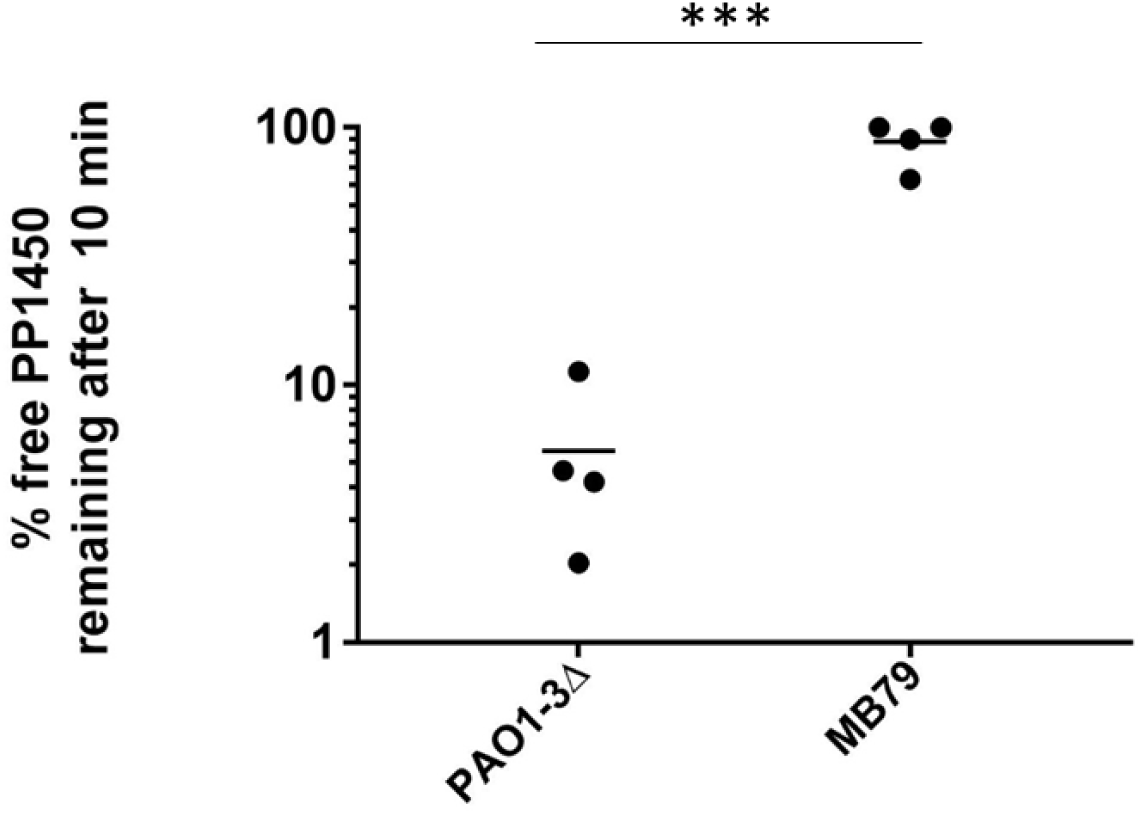
Percent of free phage remaining in the supernatant after adsorption of PP1450 for 10 minutes at 37°C in Lennox medium (*** = Unpaired T-test, P < 0.0001).

### *LS1* and *LUZ24* phage infections affect PAO1-3Δ growth differently, depending on growth conditions

The two *LUZ24* phages were able to multiply on strain PAO1-3Δ, but not sufficiently to generate visible lysis in aerated liquid medium (Lennox), nor in 96-well plates (Figure 3A) nor in large cultures (Figure 3C), and they usually did not form plaques on this medium. Therefore, we could not select any mutant resistant to *LUZ24* phages in this growth medium. We searched for conditions in which lysis would be more complete, and found that in minimal medium containing acetate as a carbon source (MMMa), PAO1-3Δ was fully lysed by the *LUZ24* phages (Figure 3B). In comparison, the two *LS1* phages were as active in rich and minimal medium (Figure 3, A and B).

**Figure 3:**
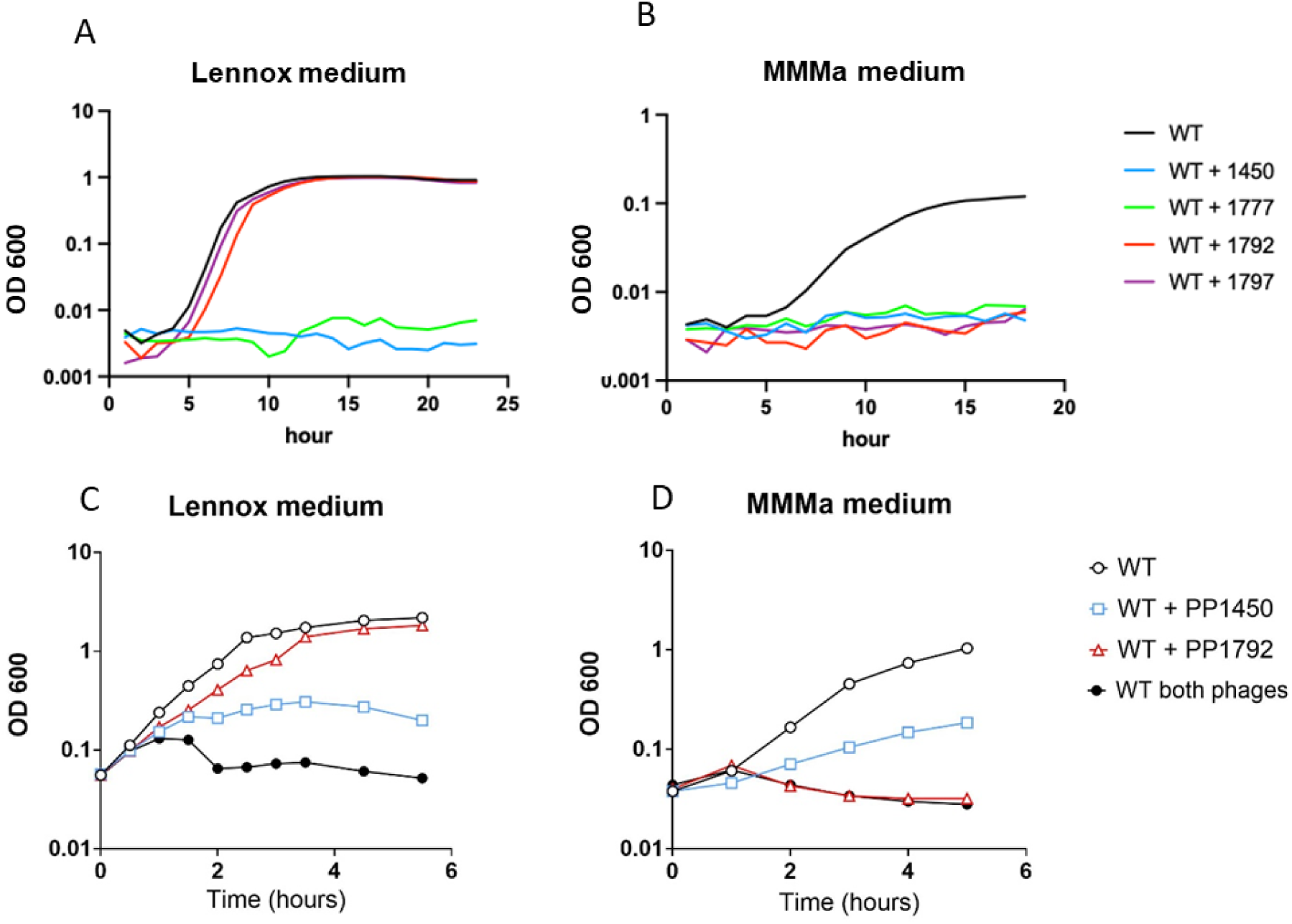
**A and B**: Growth kinetics of strain PAO1-3Δ in the presence or absence of a single phage in 96-wells plate in Lennox medium (**A**) **or** MMMa **(B)**. **C and D**: (C) PP1450 and PP1792 kinetics of lysis in large and well aerated cultures, using strain PAO1-3Δ grown in Lennox medium (C) and MMMa (D). A representative replicate of each experiment is displayed. Each experiment was repeated at least 3 times.

To further explore this improved lysis, and analyze the effect of a phage combination, the same experiment was repeated in large and aerated cultures (50 mL), using *LS1* phage PP1450, *LUZ24* phage PP1792 or a combination of both (Figure 3 C and D). In rich medium, the phage combination had a larger effect than PP1450 alone, indicating that PP1792 was not completely inactive in rich medium. In minimal medium, the reverse effect was observed, with PP1792 performing full lysis, whereas PP1450 lysis was partial, and the combination did not generate increased lysis, compared to PP1792 alone. With a different phage combination (PP1777 and PP1792), lysis by the *LS1* phage PP1777 was more pronounced than with PP1450 in aerated Lennox, and no additive effect of the *LUZ24* phage was observed (Figure S2 C).

To investigate phage multiplication in other growth media approaching conditions of *P. aeruginosa* infections in human tissues, Sputum medium, a medium mimicking the human lung sputum, and growth in partial anaerobic conditions (static Lennox supplemented with potassium nitrate, and covered with paraffin) were tested (Figure S2, A and B). In these conditions, the lytic effect of *LUZ24* phage PP1792 was again more pronounced than that of *LS1* phage PP1450. The two-phage combination had a stronger effect in Sputum medium, but not in partial anaerobic condition. Clearly, the lytic activity of the *LUZ24* phage was superior in condition of slow growth, compared to rich medium. It should be noted, however, that upon over-night incubation, bacterial growth resumed in all cases, including with the phage combination.

We conclude that the efficiency of lysis of the four phages under study depends on growth conditions, and that in some cases, the assembly of the *LS1* with the *LUZ24* phage has an additive effect.

### The polysaccharide Psl is the receptor of the two LUZ24 phages

Once conditions favorable to *LUZ24* infection were established, we used growth in MMMa medium to study the bacterial receptor of these phages. To determine at which frequency mutants resisting to *LUZ24* emerged, we conducted fluctuation assays as described above for *LS1* phages, except that Lennox was replaced by MMMa, and the plates were read after 48 of growth at 37°C. Mutation frequency was 1.8 (±1.2) ×10^-5^ per generation for PP1792, and 3.8 (±3.6) x 10^-5^ per generation for PP1797. Four mutants resistant to PP1792 were isolated. They were purified and confirmed to be P1792 resistant. To determine whether these mutants were affected at the adsorption step, adsorption efficiency after 10 minutes of incubation at 37°C was measured. Three out of the 4 mutants abolished PP1792 adsorption (Figure 4). Whole genome sequencing of these mutants revealed a mutation in the *psl* operon (Table 2): strains MB132 and MB133 had the same 13 nt deletion in *pslA*, named *pslA1*, while strain MB119 had a 17 nt deletion in *pslD*, named *pslD1*. The *pslA* gene encodes a glycosyl-transferase, while *pslD* encodes a transporter component, and both gene deletions were shown to abolish Psl production at the bacterial surface (Byrd et al., 2009). Two additional mutants obtained from the fluctuation assay with PP1797 were also sequenced, and revealed a new mutation in *pslA*, as well as a mutation in *pslH* (Table 2).

**Figure 4:**
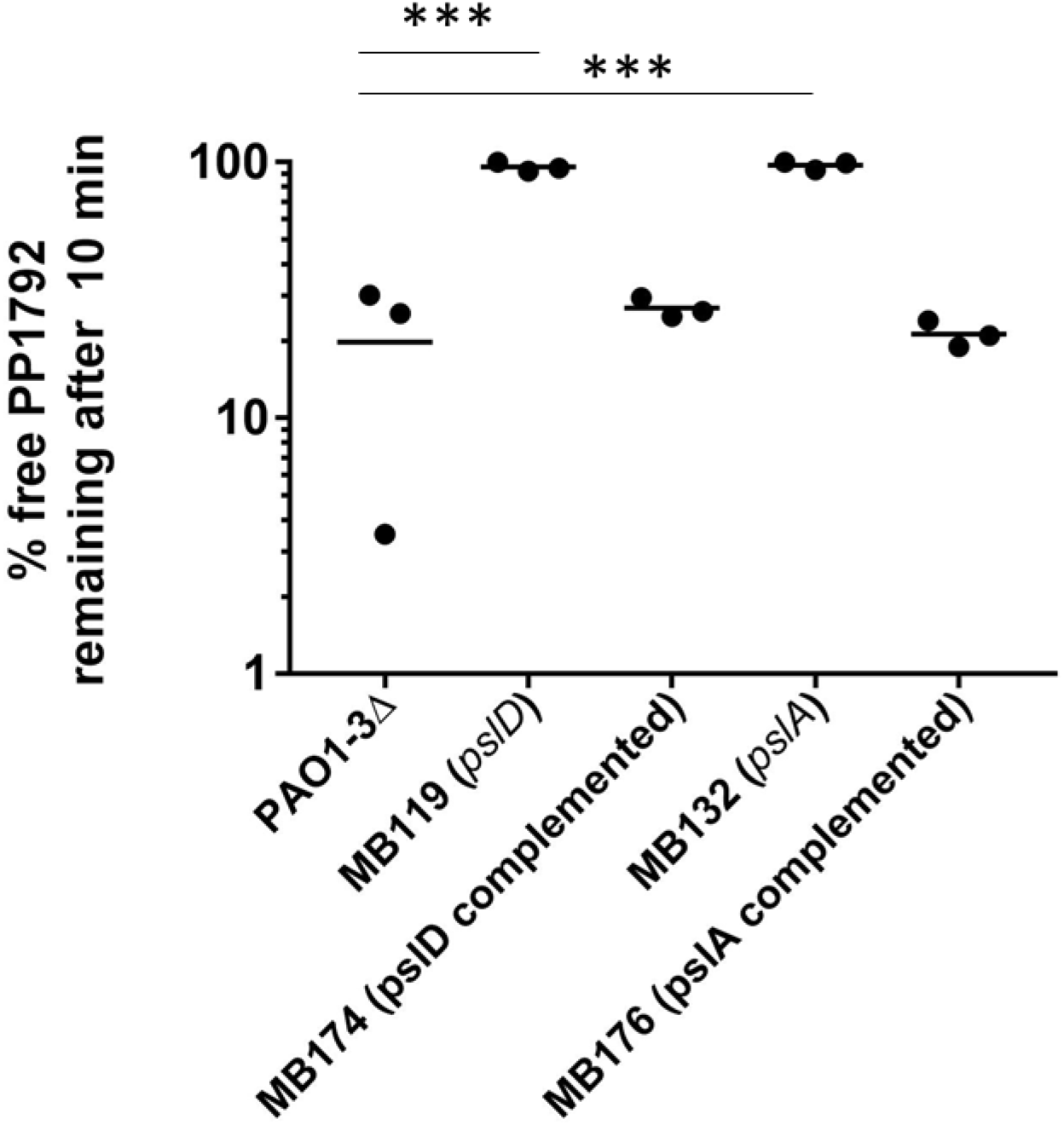
Percent of free phage remaining in the supernatant after adsorption of PP1792 for 10 minutes at 37°C in MMMa medium. One-way Anova comparisons to PAO1-3Δ, only significant values are shown (***= P < 0.001).

**Table 2:**
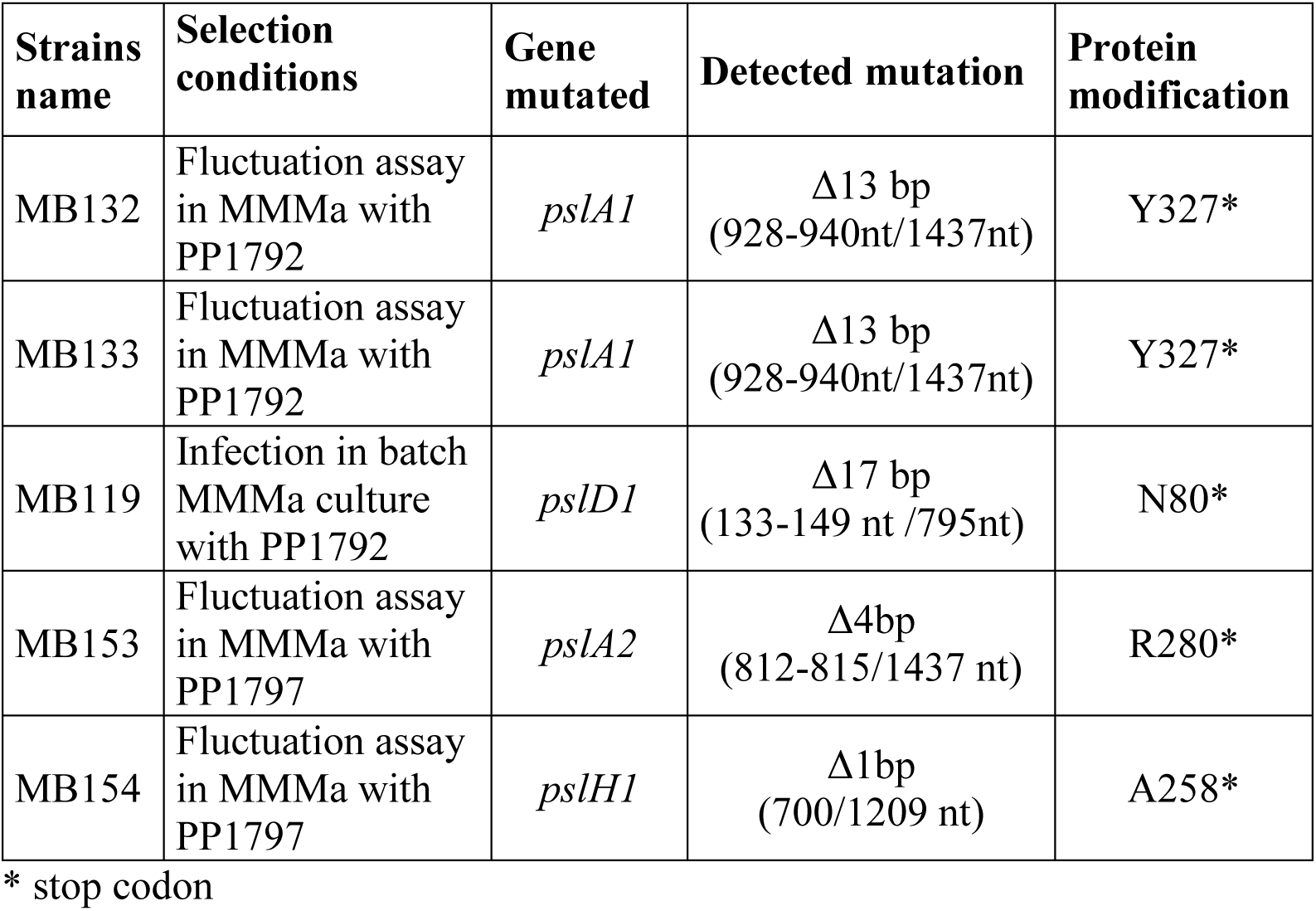
Mutations present in PAO1-3Δ adsorption mutants resisting to *LUZ24* phages.

| Strains name | Selection conditions | Gene mutated | Detected mutation | Protein modification |
| --- | --- | --- | --- | --- |
| MB132 | Fluctuation assay in MMMa with PP1792 | *pslA1* | $\Delta 13$ bp<br>(928-940nt/1437nt) | Y327* |
| MB133 | Fluctuation assay in MMMa with PP1792 | *pslA1* | $\Delta 13$ bp<br>(928-940nt/1437nt) | Y327* |
| MB119 | Infection in batch MMMa culture with PP1792 | *pslD1* | $\Delta 17$ bp<br>(133-149 nt /795nt) | N80* |
| MB153 | Fluctuation assay in MMMa with PP1797 | *pslA2* | $\Delta 4$bp<br>(812-815/1437 nt) | R280* |
| MB154 | Fluctuation assay in MMMa with PP1797 | *pslH1* | $\Delta 1$bp<br>(700/1209 nt) | A258* |

\* stop codon

To confirm that these mutations were sufficient to suppress adsorption, a complementation assay was conducted. Strains MB132 (*pslA1*) and MB119 (*pslD1*) were transformed with plasmid pPSV35-*pslA* expressing *pslA*, and plasmid pPSV35-*pslD* expressing *pslD*, respectively. These complemented strains were grown in MMMa supplemented with gentamycin, and tested for their capacity to adsorb PP1792. Phage PP1792 absorption to the *pslA* and the *pslD* complemented strains was comparable to the WT strain (Figure 4). In addition, PP1792 and PP1797 lysed the complemented strains with an efficiency of plating similar to the WT strain, while being unable to infect the *psl* mutant strains *pslA1* and *pslD1* (Table S4). All together, these data show that the receptor for the *LUZ24* phages PP1792 and PP1797 is the Psl polysaccharide.

### Psl binds more efficiently to the bacterial surface in MMMa than in Lennox, during exponential growth

The Psl polysaccharide exists both in an attached form to the bacterial surface, as well as in a free form in bacterial supernatant, and in rich medium the attached form is transiently present at the onset of the exponential phase (Melaugh et al., 2023). We therefore investigated whether the reason for the better infection of *LUZ24* phages in MMMa could be an increased fraction of bound Psl under this growth condition. Surface-bound, as well as free Psl were collected from PAO1-3Δ cultures (‘WT’ in Figure 5) grown in Lennox or MMMa, at various optical densities, and immunodetected with Psl monoclonal antibodies. We found that in MMMa, all Psl was present on the bacterial surface, and no Psl was present in the supernatant of an overnight culture. In contrast, in rich medium, surface-bound Psl was not as abundant during exponential growth, and most of Psl was found in supernatants of overnight cultures (Figure 5, Figure S3).

**Figure 5.**
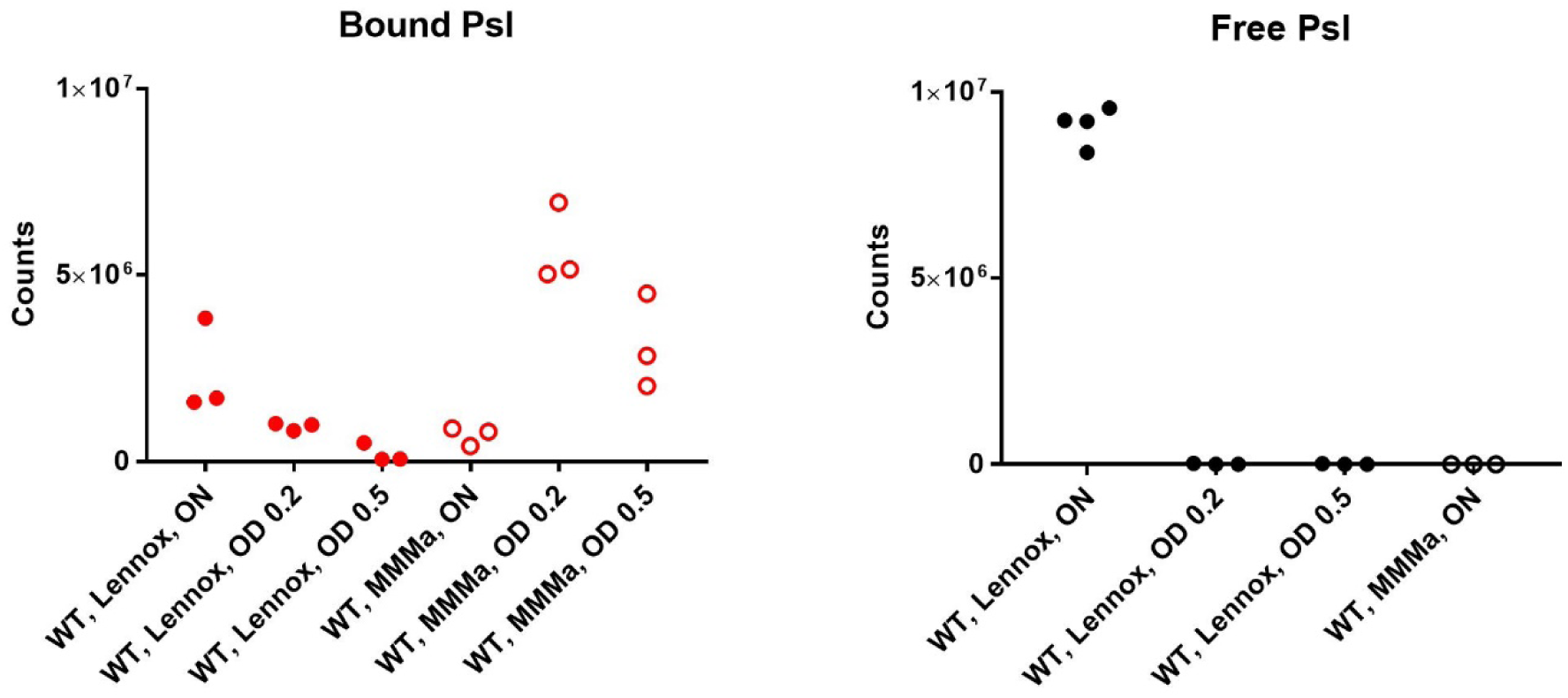
Psl amounts quantified from Western blots. Bacteria collected from over-night cultures, or during exponential growth at OD 0.2 and 0.5 were used to extract surface-bound Psl from equivalent amounts of washed pellets (left side, “bound Psl”), and these extracts were then deposited on a membrane and treated for immunodetection as described in the Methods. In parallel, 5 µL of supernatants from the same cultures (right side, “free Psl”) were directly deposited on the same membranes.

We concluded that bacteria grown in MMMa had larger amounts of bound Psl, compared to those grown in Lennox medium, especially in exponential phase, a context favorable to phage *LUZ24* multiplication.

### The Psl-binding LUZ24 phage PP1792 is more efficient at biofilm removal on intubation device than the LS1 phage PP1777

The Psl exopolysaccharide is involved in biofilm formation, which can develop on the intubation device of ventilated patients and cause VAP. To evaluate the relevance of phage therapy in such a setting, biofilm of PAO1-3Δ were grown on pieces of intubation tubes (Figure S4). After 7 h of biofilm formation at 30°C, 1.5 x 10^8^ PFU of phage were added individually or in combination (when in combination, 1.5 x 10^8^ PFU of each phage was added, so 3 x 10^8^ PFU phages in total) and incubated 17 h. The biofilm was quantified using crystal violet coloration (Figure 6A). On the PAO1-3Δ biofilm, the *LS1* phage PP1777 phage did not significantly decrease the amount of polysaccharides. In contrast, the *LUZ24* phage PP1792 significantly decreased the amount of staining of the PAO1-3Δ biofilm, by a factor of 10. The combination of the two phages significantly decreased the biofilm at level equal to that of *LUZ24* phage PP1792. The same tendency was observed for the CFU counts (Figure 6B), although differences were not significant.

**Figure 6:**
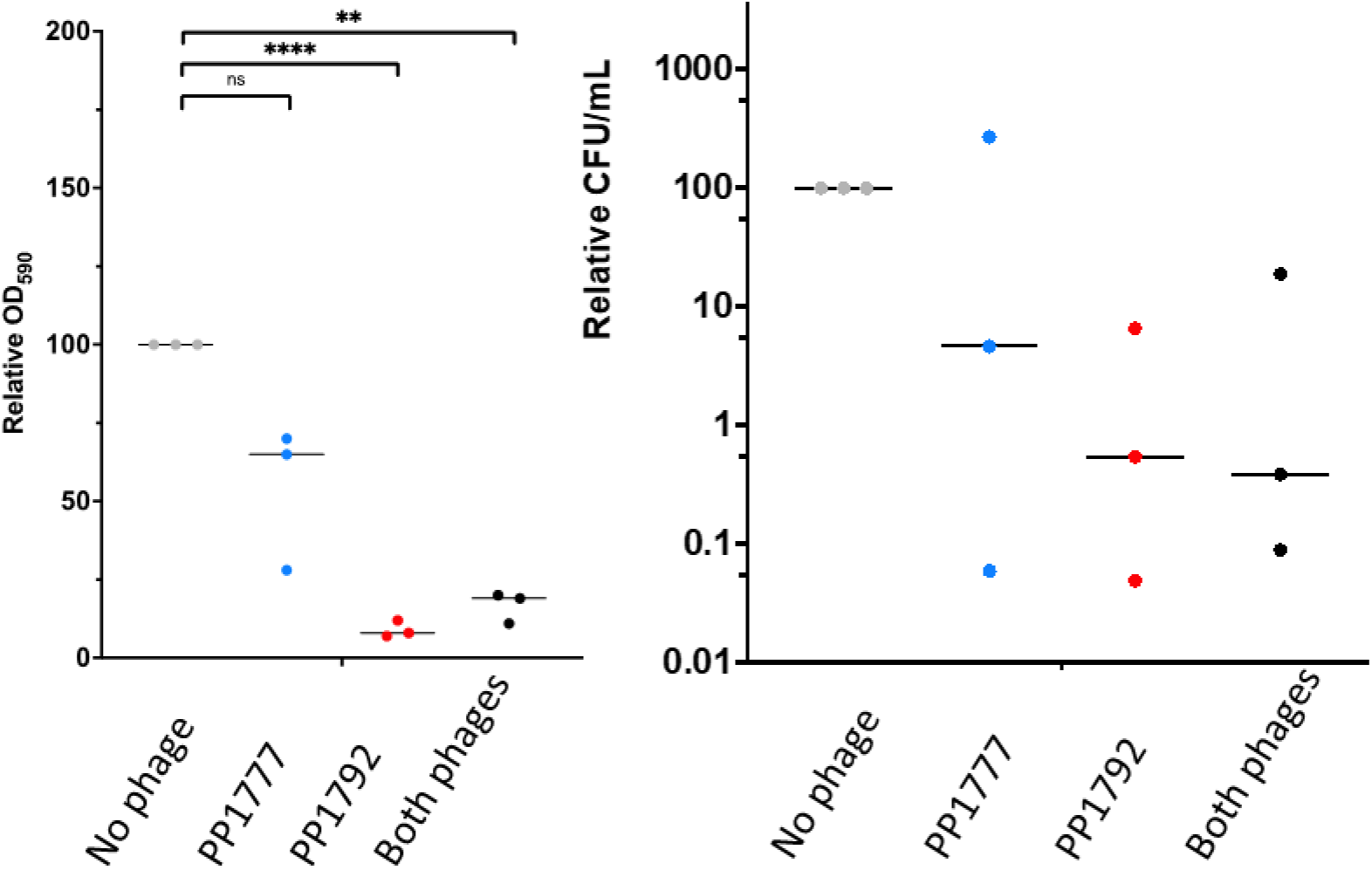
A. Biofilm quantification by crystal violet coloration, expressed as a percent of the untreated condition (grey dots). Two way Anova tests, * : P < 0.05; ** P < 0.01; **** P < 0.001; ns: non-significant. B. CFU counts, expressed relatively to the untreated condition (grey dots).

### Both LUZ24 and LS1 phages are able to kill sessile bacteria within biofilms

To explore the kinetics of phage-mediated biofilm degradation, biofilms were grown in 96-well plates, and a confocal microscopy set-up was used to track the biofilm content of living bacteria over 48 hours after phage addition. Instead of using our reference PAO1-3Δ strain, its more natural ancestor PAO1_OR, hosting two filamentous prophages was chosen. These prophages contribute to biofilm growth and stability over time (Rice et al., 2009). The strain also contained plasmid pSEVA627m, a low copy-number expressing constitutively the *gfp* gene under a strong promoter (Silva-Rocha et al., 2013), allowing to detect living bacteria.

After 6 hours of biofilm formation at 37°C, *LUZ24* phage PP1792 (5×10^6^ PFU), *LS1* phage PP1450 (5×10^6^ PFU), or a combination of both (2.5×10^6^ of each phage, 5×10^8^ total PFU) were added, and biofilm behavior was monitored. Phage addition prevented biofilm growth during the next 16 hours. Interestingly, later on, bacterial overgrowth took place for each single phage addition, but not for the phage combination (Figure S5). This indicated that phages are able to attack bacteria during the first stage of biofilm attachment, and if given in combination, to prevent overgrowth during at least 48 hours.

We next investigated whether phages can also attack a biofilm grown to maturation for 16 hours at 37°C in Lennox in a 96-well plate. After the 16 hours of static growth, 5×10^8^ PFU of a single phage, or 2.5 x 10^8^ PFU of each phage in the two-phage combination, were added, the plate was placed under the microscope at 37°C, and imaging was initiated. Three technical repeats were placed on each plate, and three biological repeats were done. One of them is shown Figure 7, and the two others are shown in Figure S6 and S7. The last image of a representative movie of each phage treatment for each biological replicate is also shown, and the corresponding movies are also available.

**Figure 7.**
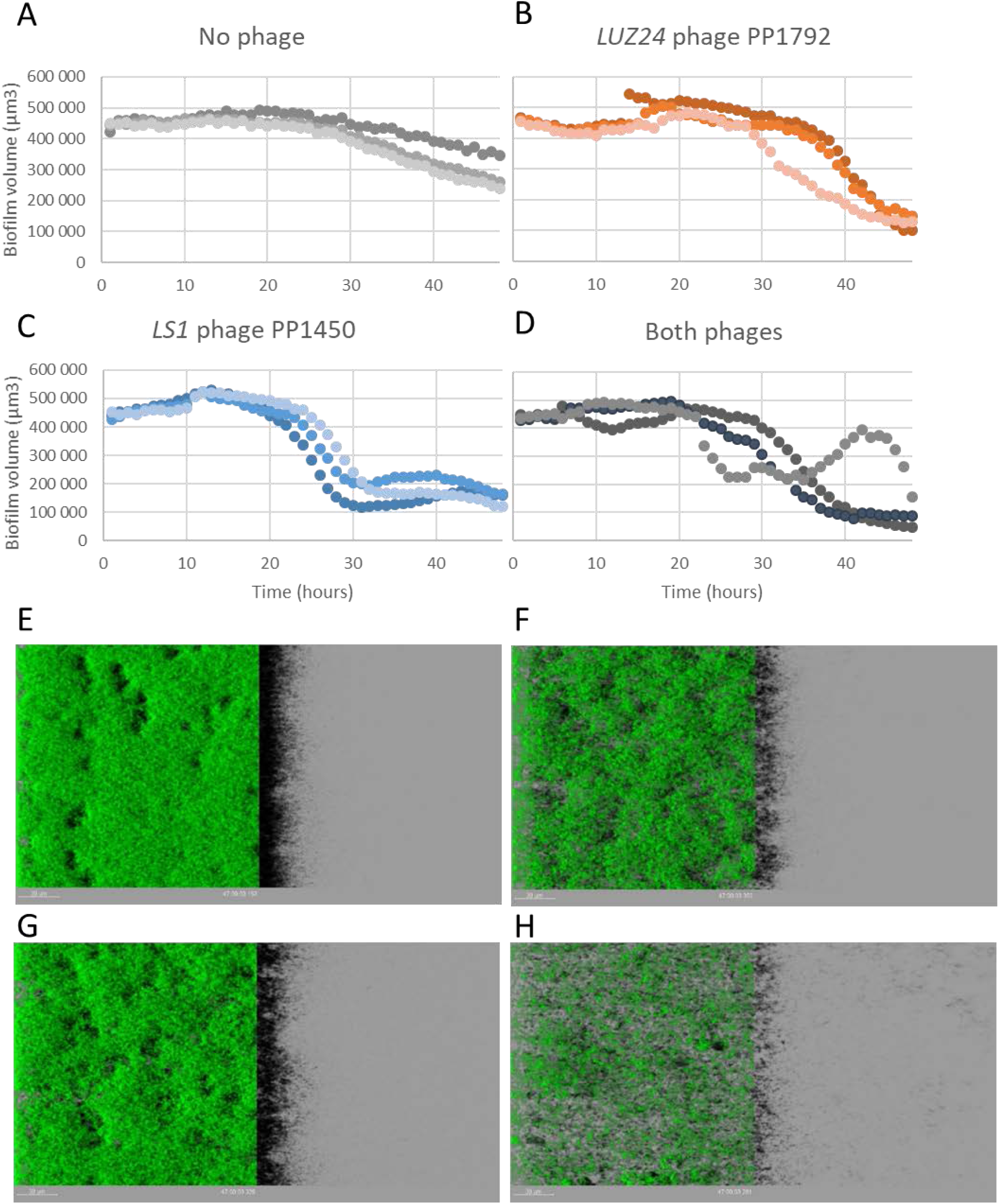
Quantification of living cell volumes (µm^3^, y axis) within a PAO1_OR biofilm, as a function of time (hours, x-axis), on a 16 hours biofilm treated with no phage (A and E), *LUZ24* PP1792 (B and F), *LS1* PP1450 (C and G), or both phages (D and H). Confocal imaging was performed every hour over 48 hours. Panels E-G show the last image of one movie by treatment.

The number of living cells in the untreated biofilm was more or less stable over the 48 hours post imaging, depending on the biological replicate (i.e. stable for 28 h Figure 7A, 48 h in Figure S6A, and 23 h in Figure S7A). The occasional late decrease in viable cells (slope of 8 000 to 14 000 µm3/h in average for the two replicates in which decay took place) was likely due to nutriment depletion over the long incubation time (64 h in total). Addition of the *LS1* phage PP1450 resulted in a steep decrease of viable cells in the biofilms (22 000 +/- 10 000 µm3/h), which always took place earlier than the natural decay, starting between 8 and 21 hours post phage addition. The *LUZ24* phage attack took place 3 to 10 hours later than *LS1* attack. In the absence of natural decay (Figure S6), lysis started 11 hours post addition (and 3 hours after *LS1*), whereas in the two replicates with natural decay, it started 2 hours later than this decay. In all cases however, a steep slope (21 000 +/- 5000 µm3/h) was observed, suggesting phage lysis was taking place.

The combination of the two phages did not manifest an additive effect in kinetics parameters (time of inflexion, slope) compared to infections with each phage alone, and followed most often the *LS1* PP1450 kinetics parameters (Figure 7D, S6D and S7D). In average, at the end point of all kinetics including the phage combination, the volume of living cells was 3.9 +/-2.2 fold less than in the t48 time point of the untreated biofilms (Figure 7 H). Interestingly, this value was twice lower than for each phage alone: the PP1792 treatment leads to a 2 +/- 1 fold final reduction in living cell volume, and this value is 1.8 +/- 1 for PP1450. This is probably due to the absence of regrowth with the phage combination, contrary to what is observed for 6 of the 17 single phage experiments (see Figure S6B and C, Figure S7B). Interestingly, one of the eight biofilms treated with the phage combination also exhibited some regrowth, but it was followed by a second lysis step (Figure 7D). Clearly, mature biofilms proved much more difficult to keep in check for the tested phages, compared to the initial stage of biofilm formation (phage addition after 6-7 hours of biofilm growth), but some signs of phage attack could be observed.

### Searching for mutants resisting to phages with different receptors: LS1 resistant strains are sensitized to LUZ24 infection in rich medium

To explore the capacity of our phage combination to prevent bacterial regrowth of resisting mutants, various phage applications to planktonic bacteria were tested. First, we examined how each phage species was growing on receptor mutants preventing the growth of the other phage species. *LS1* phage growth was effective on *pslA* and *pslH* mutants, and *LUZ24* phage growth was effective as well on the *wzy1* mutant strain MB79, on MMMa. Interestingly, on this MB79 mutant strain, *LUZ24* growth became effective even in rich medium (Figure S8 and Table S3).

Searching for the reason of such an improvement, we first investigated whether surface-bound Psl was increased for this mutant, in rich medium. This was not the case (Figure S9). We next investigated whether adsorption efficiency was improved, and found that *LUZ24* phage PP1792 adsorption was significantly more efficient on strain MB79 (*wzy1*), compared to PAO1-3Δ in Lennox (Figure 8A). The fact that phage adsorption is more efficient, while receptor concentration is not increased, suggests that the default of *LUZ24* infection in rich medium is due to a shielding effect of the O-antigen chain of the LPS.

**Figure 8.**
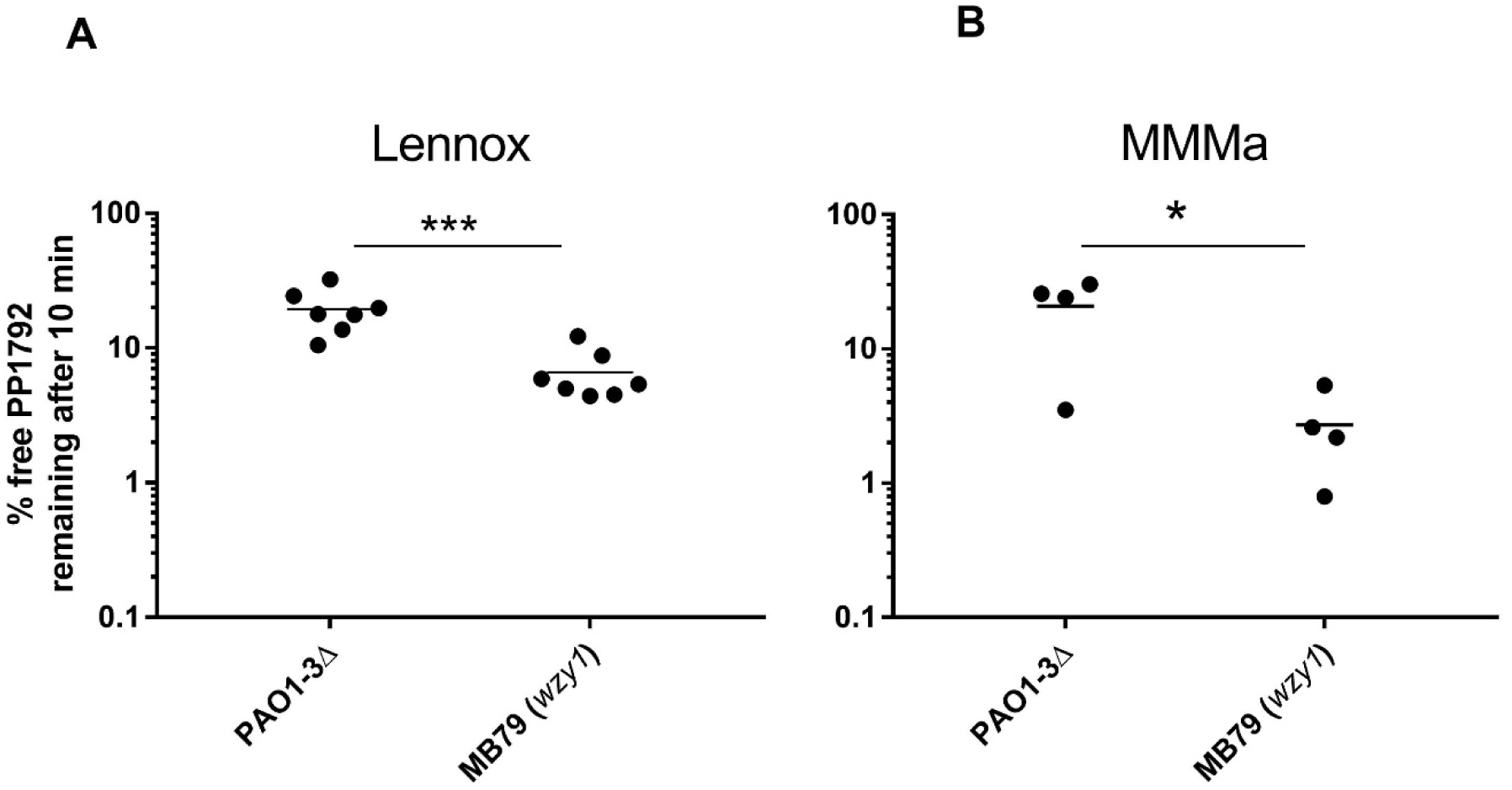
Percent of free phage remaining in the supernatant after adsorption of PP1792 for 10 minutes at 37°C on PAO1-3Δ and its *wzy1* derivative MB79, in rich Lennox medium (A) and in MMMa minimal medium (B). Unpaired T-test comparisons (Student test, ***: P=0.1%, *: P = 2%)

Interestingly, in minimal medium, *LUZ24* phages were able to infect the PAO1-3Δ, despite the presence of the O-antigen chain (Figure 3B and D), suggesting that this shielding effect was overcome. This may be due to the more elevated amounts of surface-bound Psl, in this medium, permitting to counterbalance the O-antigen shielding effect. The two effects are probably additive, as we observed that *LUZ24* phage PP1792 adsorption was also more efficient on the *wzy1* mutant than on PAO1-3Δ, in minimal medium (Figure 8B). This shielding effect was also visible by eye: in the presence of the O-antigen chain, *LUZ24* phages formed turbid plaques, on MMMa as well as on Lennox (on this medium, some plaques were sporadically detected), whereas they formed clear plaques on the *wzy1* mutant strain MB79 (both in Lennox and MMMa). An interesting consequence of these analyses is that the O-antigen mutants that are often selected by *LS1* phage infection, are sensitized to *LUZ24* phage infections in rich medium.

To deepen our understanding of the emergence of phage resisting mutants, we next analyzed a set of five independent mutants emerging from PAO1-3Δ co-infected with *LS1* phage PP1777 and *LUZ24* phage PP1792, upon growth in Lennox liquid cultures. These mutants were resistant to all four phages (Table S5). Whole genome sequencing revealed for 3 of them a mutation or a deletion involving the *galU* gene, which function is needed for both LPS and Psl synthesis. The fourth mutant was in gene *PAO1OR_5105* of unannotated function. InterPro annotates the corresponding protein (accession K9HUG4) as a putative glycosyl transferase involved in LPS synthesis. The last mutant affected the *ygiF* gene, encoding a putative inorganic triphosphatase. While speculative, it is possible that YgiF influences the synthesis of LPS and Psl. By hydrolyzing inorganic triphosphate, YgiF may help maintain calcium homeostasis (Kohn et al., 2012), which is crucial for regulating polysaccharide production. Calcium ions play a key role in biofilm formation, and disruptions in calcium signaling can impact the synthesis of LPS and Psl, particularly in species like *P. aeruginosa* (Kolodkin-Gal et al., 2023; Pazol et al., 2022).

Finally, to mimic successive infection by an *LS1* phage followed by a *LUZ24* phage, we searched for resistant mutants, starting from the *wzy1* mutant strain MB79, after infection with *LUZ24* phage PP1792 and overgrowth in liquid Lennox medium. Five mutants were purified and analyzed for their sensitivity profile to the four phages (Table S6). Most strains (4/5) had the same phenotype, they were totally resistant to the two *LUZ24* phages and remained resistant to *LS1* phage PP1450, but became partially sensitive to *LS1* phage PP1777. The last mutant, MB99, was different: it was totally resisting to the two *LS1* phages, and to *LUZ24* phage PP1797, but remained sensitive to PP1792. These mutants were totally sequenced. Four of them had a mutation in the *fklB* gene, coding for a peptidyl-prolyl cis-trans isomerase (PPiase) of the FkpA family. FkpA is a periplasmic protein having both a chaperone and a PPiase function in *Escherichia coli* (Cumby et al., 2015). The last mutant MB99 had a different mutation, in the *rocS1* gene (Table S6). Adsorption assays of the *LUZ24* phage PP1792 were conducted on strains MB91 and MB92, which are two different *wzy fklB* double mutants, as well as on MB99, the *wzy rocS1* mutant (Figure S10). They revealed that the *fklB* mutation did not affect the adsorption step, while the *rocS1* mutation did.

Interestingly, none of these 5 mutants had a mutation in either the *pslA* or the *plsD* gene, suggesting that the double *wzy psl* mutant might not be viable, at least in rich medium. The search for resisting mutants was therefore repeated in MMMa, and four resistant clones were purified and analyzed for their sensitivity profiles. Two had a typical *galU* profile (resisting to all phages), and the last two were resistant to the two *LUZ24* phages, but sensitive to the two *LS1* phages (MB187 and MB189, Table S7). These two strains were sequenced, they had three mutations: the initial *wzy1* mutation of strain MB79, a mutation in either *pslA* or *pslH,* and a last mutation in *wbpL*, adding a G in a run of 9Gs. This gene codes for a glycosyl transferase needed for the synthesis of both A and B chains of the O-antigen.

## DISCUSSION

In this study, we investigated the phage-host relationships between a prophage-less derivative of the PAO1_OR strain of *P. aeruginosa*, PAO1-3Δ, and four phages selected for their capacity to kill collectively the 43 strains of an international reference panel (De Soyza et al., 2013). Two of these phages belonged to the *Pbunavirus LS1* species, and we determined that their receptor was the O-antigen of LPS, as expected for this species. The two others belonged to the *Bruynoghevirus LUZ24* species, and for these phages, although characterized long ago (Ceyssens et al., 2008), the host receptor remained unknown. We were able to find the receptor of the *LUZ24* phages PP1792 and PP1797, by playing with the bacterial growth conditions of PAO1-3Δ. Whereas inefficient lysis of PAO1-3Δ was observed in Lennox medium liquid culture, and only turbid spots at high phage concentrations were visible on Lennox agar plates, the phages inhibited growth in liquid in minimal medium MMMa, and formed individual turbid plaques on this medium. We also observed that *LUZ24* infection was more efficient upon bacterial growth in Sputum medium, as well as under partial anaerobic conditions (i.e. upon static growth with a paraffin top layer). These *LUZ24* phages should therefore be active when their host has a metabolism adapted to slow growth, such as inside human wounds or lungs.

Efficient *LUZ24* phage growth in MMMa allowed for the selection of bacterial mutants resisting to phage infection, and 5/6 of these were adsorption mutants. We determined that *LUZ24* phages use the polysaccharide Psl to bind to the bacterial surface. We obtained *psl* mutants with altered *pslA, pslH* or *pslD* genes, all of which are necessary for the synthesis of the Psl polysaccharide (Byrd et al., 2009). Recently, an independent study reached the same conclusion that Psl was the primary receptor of their *LUZ24* phages, using a completely different approach (Walton et al., 2024). In line with our results, we noted that among mutants selected with still another *Bruynoghevirus* phage, named FJK, a *pslA* mutant was obtained, among others (Kunisch et al., 2024).

We also found that Psl was more abundant at the bacterial surface of exponentially growing cultures in minimal, MMMa medium, compared to rich, Lennox medium. This may explain why MMMa is a favorable context for phage growth, since the receptor is permanently available at the surface. Although overnight culture in rich medium contained large amounts of free Psl, we did not get evidence that this free Psl was competing for phage adsorption, since inoculation of rich medium with washed bacteria did not allow to improve bacterial lysis in this medium.

Since Psl is the main polysaccharide forming the biofilm matrix in strain PAO1, we tested the phage efficiencies on PAO1-3Δ biofilms formed during 7 hours on the tubing of an intubation device. The biofilm matrix was diminished by a factor of 10 upon incubation for 16 hours with *LUZ24* phage PP1792, whereas it was not significantly affected by *LS1* phage PP1777 treatment at this time point. We next tested phage activity on mature biofilms grown statically on 96-well plates (16 hours of growth before infection), using the parental, PAO1_OR strain, which was expected to be more difficult to eradicate, because its prophages contribute to biofilm stability over time (Rice et al., 2009). Moderate lysis occurred with both single phage treatments and started sooner with the *LS1* phage PP1450. Some regrowth was observed for 4 of the 9 repeats with *LUZ24* phage PP1792, and 2 of the 8 repeats with *LS1*. The combination of the two phages did not lead to a more rapid killing of the biofilm, compared to the more rapid phage alone, but interestingly, no visible overgrowth was observed for the 8 replicates, during the 48 hours of the assay. This contrasts with an earlier report, where phages belonging to the same genera, even when applied in combination to a 24 h biofilm of PAO1, led to biofilm regrowth (measured with crystal violet) 8 hours post phage treatment (Pires et al., 2017). We were surprised by the slow kinetics of phage attack on mature biofilms, and cannot exclude that the peptidoglycan released upon phage-mediated bacterial lysis stimulates biofilm formation, as recently observed for various bacterial species, including *P. aeruginosa* (Vaidya et al., 2025).

The complementary lytic characteristics of the *LS1* phages and the *LUZ24* phages could be beneficial in the frame of phage therapy. While the pair of *LS1* phages multiplied more efficiently on bacteria grown in aerated Lennox medium, the *LUZ24* phage pair had the opposite behavior, multiplying more efficiently on bacteria grown in minimal medium, as well as in Sputum medium or upon limited oxygen availability. On an infection site, there might be bacteria in different physiological states and by combining these two phages, more bacteria should be killed, whichever state they are in.

As our two phage species targeted different receptors, their simultaneous application was expected to limit the emergence of resistant mutants (Hesse et al., 2020; Yang et al., 2020). We found, however, that *galU* mutants were resisting to both phage species, so that the gain provided by the phage combination was not as high as expected. Nevertheless, in our screenings *galU* deletions arose at a lower frequency compared to *wzy* mutations (only 1/10 of *LS1* resisting mutants were *galU* deletions). Moreover, in our plate assay, biofilm overgrowth did not appear over 48 h when a phage combination was applied to early (6 h) or late (16 h) biofilms, suggesting again that such mutants are rare. Unexpectedly, *wzy* mutants resisting to *LS1* phages (which occur 10 times more frequently than *galU* mutants) were sensitized to the *LUZ24* phages, in the rich medium where normally *LUZ24* phages are less active. This is an advantage of this phage combination.

The impact of the culture medium on phage-host interaction has not been thoroughly investigated until now, although it starts being considered. Lourenço et al. observed that in the gut microbiota, LPS host receptors on the *Escherichia coli* surface are not produced at sufficient levels to permit phage CLB-P1 multiplication (Lourenco et al., 2022). Moreover, the N4 phage receptor, an *E. coli* phage known for a long time, was also recently identified, thanks to the appropriate choice of culture conditions (Sellner et al., 2021). This receptor is a novel surface glycan named NGR, which existence had escaped scrutiny until now. Interestingly, synthesis of this receptor was dependent on the concentration of cyclic di-GMP in the cytoplasm, which itself was influenced by the levels of enzymes needed to synthetize or degrade cyclic di-GMP.

In one of our genetic screens aiming at characterizing bacterial mutant strains able to overcome phage infection, we searched for mutants arising from the successive application of a *LS1* phage (which generated mostly *wzy* mutants) followed by application of a *LUZ24* phage, in rich medium. The screen identified a single adsorption mutant, which had a *rocS1* mutation. This gene, also known as *sadS*, is a histidine kinase belonging to an intricate “two-component system” made of two sensors (*rocS1* and *rocS2*) and three regulators (*rocA1*, *rocA2* and *rocR*). While RocA1 and RocA2 are HTH-containing DNA binding proteins, the RocR regulator is different: it contains a diesterase domain, and decreases the levels of cyclic di-GMP (Rao et al., 2008). By analogy with the N4 observations, we therefore hypothesize that a way to resist *LUZ24* phages is to modify the intracellular level of cyclic di-GMP, which might indirectly command the level of synthesis of the Psl polysaccharide, and thereby alter as well biofilm formation.

All other mutants characterized in this screen mimicking successive application were not adsorption mutant, and had a mutated FklB. Two functions related to phage cycles are reported in the literature for FklB homologs. The FklB protein is homologous to FkpA and SlyD proteins of *E. coli*. FkpA is a periplasmic protein necessary for HK97 phage injection (Cumby et al., 2015). SlyD facilitates lysis during the lytic cycle phage ΦX174, by stabilizing its protein E, an inhibitor of MraY (Bernhardt et al., 2002). These two functions would enter into play after the initial adsorption step, and are compatible with defect posterior to adsorption, as we observed for the *fklB* mutants.

Overall, this genetic identification of the *LUZ24* phage receptor Psl allowed to uncover important characteristics of its infection process, namely its dependence on bacterial growth conditions. We also demonstrate here that the combination of the four phages selected for compassionate phage therapy treatments, two *LUZ24* and two *LS1* phages, have differing and complementary properties making them an appropriate combination for treatments.

## Acknowledgements.

We would like to thank Isabelle Vallet for providing plasmids and for her constant help at the onset of the project with *Pseudomonas aeruginosa* genetics, Scott Rice for the gift of strain PAO1ΔPf4, and Daniel Wozniak for his advice at the onset of Psl immunodetection experiments. The members of the Migale platform are also thanked for the bio-informatics tools made available and their constant support.

The authors declare they have no competing interests.

## Notes

### Competing Interest Statement

The authors have declared no competing interest.

