## Supplemental tables and figures for "Complementary killing activities of *Pbunavirus LS1* and *Bruynoghevirus LUZ24* phages on planktonic and sessile *Pseudomonas aeruginosa* PAO1 derivatives"

### Supplementary material

Table S1. Bacteria and plasmids used in this study

| Strain or plasmid | Genotype and/ or relevant characteristics | Source or reference |
|---|---|---|
| *P. aeruginosa PAO1 derivatives* | | |
| PAO1_OR | | LN871187 |
| PAO1-3Δ | PAO1-OR ΔPf4::FRT Δ*rep*_Pf6::FRT Δ*toxA* | This study |
| MB75 | PAO1-3Δ *wzy2 gloA2* | This study |
| MB76 | PAO1-3Δ *wzy3* | This study |
| MB078 | PAO1-3Δ with 329 240 bp deletion | This study |
| MB079 | PAO1-3Δ *wzy1* | This study |
| MB132 | PAO1-3Δ *pslA1* | This study |
| MB133 | PAO1-3Δ *pslA1* | This study |
| MB119 | PAO1-3Δ *pslD1* | This study |
| MB153 | PAO1-3Δ *pslA2* | This study |
| MB154 | PAO1-3Δ *pslH1* | This study |
| MB191 | PAO1-3Δ pSEVA627M | This study |
| MB198 | PAO1-12228 pSEVA627M | This study |
| CRR105 | PAO1-3Δ Δ(*PAO1OR_2949 –2986*) | This study |
| CRR107 | PAO1-3Δ Δ(*PAO1OR_2949 –2986*) | This study |
| CRV105 | PAO1-3Δ *galU* | This study |
| CRV109 | PAO1-3Δ *PAO1OR_5105* | This study |
| CMR105 | PAO1-3Δ *ygiF1* | This study |
| MB087 | PAO1-3Δ *wzy1 fklB1* | This study |
| MB096 | PAO1-3Δ *wzy1 fklB2* | This study |
| MB091 | PAO1-3Δ *wzy1 fklB1* | This study |
| MB092 | PAO1-3Δ *wzy1 fklB2* | This study |
| MB099 | PAO1-3Δ *wzy1 rocS1* | This study |
| MB187 | PAO1-3Δ *wzy1 wbpL pslH* | This study |
| MB189 | PAO1-3Δ *wzy1 wbpL pslA* | This study |
| *E. coli strains* | | |
| β2163 | Diaminopurine dependent growth, RP4 conjugation genes inserted into the chromosome | (Demarre et al., 2005) |

| | | |
|---|---|---|
| JM105 | *endA*1 *glnV*44 *sbcB*15 *rpsL thi*-1 Δ(*lac-proAB*) [F' *traD36 proAB*+ *lacIq lacZ*ΔM15] *hsdR*4(rK-mK+) | (Yanisch-Perron et al., 1985) |
| *Plasmids* | | |
| pPSV35 | Shuttle vector with gentamicin resistance gene (*aacCA1*), PA origin, *lacIq*, and *lacUV5* promoter | (Rietsch et al., 2005) |
| pSEVA627m | Shuttle vector with a msfGFP, RK2 origin, and gentamicin resistance gene | (Silva-Rocha et al., 2013) |
| pPSVA | *pslA* gene from PAO1-3Δ cloned into pPSV35 | This study |
| pPSVD | *pslD* gene from PAO1-3Δ cloned into pPSV35 | This study |
| pEX18ApGW | Not replicative in *P. aeruginosa*, contains a *sacB* gene and the *oriT* origin. | (Choi and Schweizer, 2005) Accession number AY028469 |
| pEXG2 | Not replicative in *P. aeruginosa*, contains a *sacB* gene and an *aac1* gene that is not flanked by FRT sequences. | Accession number KM887143 |
| pFlp2 | Contains the flippase gene | Accession number AF048702 |
| pMB01 | Integration between *Kpn*I and *Hind*III of pEX18ApGW of a 1660 bp fragment encompassing the PAO1 sequences flanking Pf4, with Pf4 itself being replaced by FRT-*aac1*-FRT. This fragment was amplified on strain PAO1ΔPf4 from (Rice et al., 2009). Left and right flanks are delimited by oligonucleotides (Pf4_BG_5' and 3') and (Pf4BD_3' and 5') | This study |
| pMB03 | Integration between *Kpn*I and *Hind*III of pEX18ApGW of a 1850 bp fragment encompassing the PAO1 sequences flanking the *rep* gene of Pf6, with *rep* itself being replaced by FRT-*aacA1*-FRT. Left and right flanks are delimited by oligonucleotides (Pf6_rep_BG_5' and 3') and (Pf6_rep_BD_3' and 5') | This study |
| pMB06 | Integration between *Eco*RI and *Hind*III of pEXG2 of a 676 bp fragment encompassing the PAO1 sequences flanking the *toxA* gene, placed directly next to each other. Left and right flanks are delimited by oligonucleotides (exoA_BG_5' and 3') and (exoA_BD_3' and 5') | This study |

Table S2: Primers used in this study

| Name | Description | Sequence |
|---|---|---|
| **GmF** | forward primer for the amplification of the gentamicin cassette | CGAATTAGCTTCAAAAGCGCTCTGA |
| **GmR** | reverse primer for the amplification of the gentamicin cassette | CGAATTGGGGATCTTGAAGTTCCT |
| **Pf4_BG_5'** | forward primer for the amplification of Pf4 left border (restriction enzyme: KpnI) | CATGGTACCTGGCAGCAGACCCAGGACGC |
| **Pf4_BG_3'** | reverse primer for the amplification of Pf4 left border | AGGAACTTCAAGATCCCCAATTCGCGTCA TGAGCTTGGGAAGCT |
| **Pf4_BD_5'** | forward primer for the amplification of Pf4 right border | TCAGAGCGCTTTTGAAGCTAATTCGGATC CCAATGCAAAAGCCCC |
| **Pf4_BD_3'** | reverse primer for the amplification of Pf4 right border (restriction enzyme: HindIII) | CATAAGCTTTCTGGGAATACGACGGGGGC |
| **Pf6_rep_BG_5'** | forward primer for the amplification of the left border of Pf6 *rep* gene (restriction enzyme: KpnI) | CATGGTACCAGTAGCGTTGCTCGTGATTG |
| **Pf6_rep_BG_3'** | reverse primer for the amplification of the left border of Pf6 *rep* gene | AGGAACTTCAAGATCCCCAATTCGGGC GATATGACGGTACGCAAG |
| **Pf6_rep_BD_5'** | forward primer for the amplification of the right border of Pf6 *rep* gene | TCAGAGCGCTTTTGAAGCTAATTCGATCAT TTGTGCCTCCAG |
| **Pf6_rep_BD_3'** | reverse primer for the amplification of the right border of Pf6 *rep* gene (restriction enzyme: HindIII) | CATAAGCTTGCTGCGCCTGTTGATCTG |
| **exoA_BG_5'** | forward primer for the amplification of the left border of *exoA* (restriction enzyme: HindIII) | CATAAGCTTCTCGATAGTCGCCAGGACAC |
| **exoA_BG_3'** | reverse primer for the amplification of the left border of *exoA* (restriction enzyme: BamHI) | CATGGATCCGAAGGATGAGGCTGATCGAG |
| **exoA_BD_5'** | forward primer for the amplification of the right border of *exoA* (restriction enzyme: BamHI) | CATGGATCCATCCTGATAGGTAATCCGC |
| **exoA_BD_3'** | reverse primer for the amplification of the right border of *exoA* | CATGGAATTCGGACATCACCGAGTAGATGC |

| | | |
|---|---|---|
| | (restriction enzyme: EcoRI) | |
| **pslA_5'** | forward primer for amplification of pslA | TCTGGTACCGCGTTCATCAGTAGACTTCC |
| **pslA_3'** | reverse primer for the amplification of pslA | TCTGAATTCTCGGCAGAGCAAACAAC |
| **pslD_5'** | forward primer for amplification of pslD | TCAGGTACCGGCGATGTCGCTCCTCAG |
| **pslD_3'** | reverse primer for the amplification of pslD | TCAGAATTCCTGGAAGTGAACATCATGAC |

Table S3: Efficiency of plating of the four phages on PAO1-3Δ and its *LS1* resistant derivatives, in Lennox. (+) = EOP > 10%; (+/-) = EOP between 10% and 1%; (-/+) = EOP between 1% and 0.01%; (-) = EOP < $10^{-5}$.

| **Strain** | **Efficiency of plating on Lennox** | | | | **Locus mutated** (PAO1_OR id) | **Nt or amino-acid changes** |
|---|---|---|---|---|---|---|
| | **PP1450** | **PP1777** | **PP1792** | **PP1797** | | |
| PAO1-3Δ | + | + | - | - | | |
| MB79 | - | -/+ | + | + | *wzy* (1817) | T54* |
| MB81 | - | -/+ | + | -/+ | *wzy* (1817) | R74* |
| MB75 | - | -/+ | - | - | *wzy* (1817), *gloA2* (4299) | Wzy_R74* GloA2_R50H |
| MB78 | -/+ | - | - | - | Δ(2768-3055) | 329 240 bp deletion (includes *galU*) |

Table S4: Efficiency of plating of *LUZ24* phages PP1792 and PP1797 on *psl* mutants and complemented strains.

| Bacterial strain, mutation | Complementation plasmid | Efficiency of plating | |
|---|---|---|---|
| | | PP1792 | PP1797 |
| PAO1-3Δ, WT | pPSV35 | 1 | 1 |
| MB119, *pslD1* | pPSV35 | <10-5 | <10-5 |
| MB119, *pslD1* | pPSV-D | 1 | 1 |
| MB132, *pslA1* | pPSV35 | <10-5 | <10-5 |
| MB132, *pslA1* | pPSV-A | 1 | 1 |

Table S5. Phage resistant mutants isolated from a co-infection with *LS1* phage PP1777 and *LUZ24* phage PP1792 in Lennox (strains CRR and CRV), or in MMMa (strain CMR).

| Bacterial strain | **Efficiency of plating of** | | | | **Locus mutated** (PAO1_OR id) | **Nt or amino acid changes** |
|---|---|---|---|---|---|---|
| | **PP1450** | **PP1777** | **PP1792** | **PP1797** | | |
| CRR105 | - | - | - | - | Δ(2949 –2986) | 40224 bp deletion (includes *galU*) |
| CRR107 | - | - | - | - | Δ(2949 –2986) | 40224 bp deletion (includes *galU*) |
| CRV105 | - | - | - | - | *galU1* (2967) | D130Y |
| CRV109 | - | - | - | - | (5105) | frameshifted protein |
| CMR105 | - | - | - | - | *ygiF1* (5315) | N29H |

Table S6: Efficiency of plating of the four phages on mutants resisting to PP1792, starting from strain MB79 (*wzy1*). Mutants were selected either in Lennox (MB87 to MB99) or in MMMa (MB187 and MB189). (+) = EOP > 10%; (+/-) = EOP between 10% and 1%; (-/+) = EOP between 1% and 0.01%; (-) = EOP < $10^{-5}$.

| Bacterial strain | Efficiency of plating of | | | | Additional locus mutated, relative to MB79 (PAO1_OR id) | amino acid changes |
|---|---|---|---|---|---|---|
| | **PP1450** | **PP1777** | **PP1792** | **PP1797** | | |
| MB87 | - | +/- | - | - | *fklB1* (4644) | Q74* |
| MB96 | - | +/- | - | - | *fklB2* (4644) | V13D |
| MB91 | - | +/- | - | - | *fklB1* (4644) | Q74* |
| MB92 | - | +/- | - | - | *fklB2* (4644) | V13D |
| MB99 | - | - | +/- | - | *rocS1* (1016) | P724L |
| MB187 | + | + | - | - | *wbpL* (1886)*, *pslH* (2750) | frameshifted PslH |
| MB189 | + | + | - | - | *wbpL* (1886)*, *pslA* (2757) | PslA_R297* |

*Same mutation in the two clones, WbpL_G92*

Figure S1: polymorphism of the tail gene (ORF58) of phages PP1792, PP1797 and Luz24. Diverging amino-acids of phage PP1797, relative to PP1792 and Luz24 are shown in red.

```
1797     MGLEVATYINQLVPTNPTGSDLKSFGDDHLRLIKSAIKNTFPNISQAVTVTAAQLNAVAD  60
LUZ24    MGLEVATYINQLVPTNPTGSDLKSFGDDHLRLIKSAIKNTFPNISQAVTVTAAQLNAVAD  60
1792     MGLEVATYINQLVPTNPTGSDLKSFGDDHLRLIKSAIKNTFPNISQAVTVTAAQLNAVAD  60

1797     TTQYVKPGMVIMWAGSLAQIPAGWKLCNGVGTTSNGIPVPNLIGAFPWGIDGSSQAVGTR  120
LUZ24    TTQYVKPGMVIMWAGSLAQIPAGWKLCNGVGTTSNGIPVPNLIGAFPWGIDGSSQAVGTR  120
1792     TTQYVKPGMVIMWAGSLAQIPAGWKLCNGVGTTSNGVPVPNLIGAFPWGIDGSSQAVGTR  120

1797     GGSANIVWDGFTEGTALTLAQIPSHTHTWRTRGAAALVGSAGDSGALTGGSGNAANTNLE  180
LUZ24    GGSANIVWDGFTEGTALTLAQIPSHTHTWRSRGATTLTGSAGDSGALTGGSGNAANTNLE  180
1792     GGSANIVWDGFTEGTALTLAQIPSHTHTWRSRGATTLTGSAGDSGALTGGSGNAANTNLE  180

1797     TGPAGQGQTHNHAVKINMPLGNIPPFCSVFFIIKN  215
LUZ24    TGPAGQGQTHNHAVKINMPLGNIPPFCSVFFIIKN  215
1792     TGPAGQGQTHNHAVKINMPLGNIPPFCSVFFIIKN  215
```

Figure S2: **A and B**: *LS1* phage PP1450 and *LUZ24* phage PP1792 kinetics of lysis in 40 mL cultures of PAO1-3Δ grown statically, with a paraffin layer to limit O2 diffusion inside the culture (A) or Sputum medium (B)**. C:** Comparison of lytic activities of *LS1* phage PP1777 and *LUZ24* phage PP1792 upon infection of PAO1-3Δ in 40 mL aerated Lennox cultures. A representative replicate of each experiment is displayed. Each experiment was repeated at least twice.

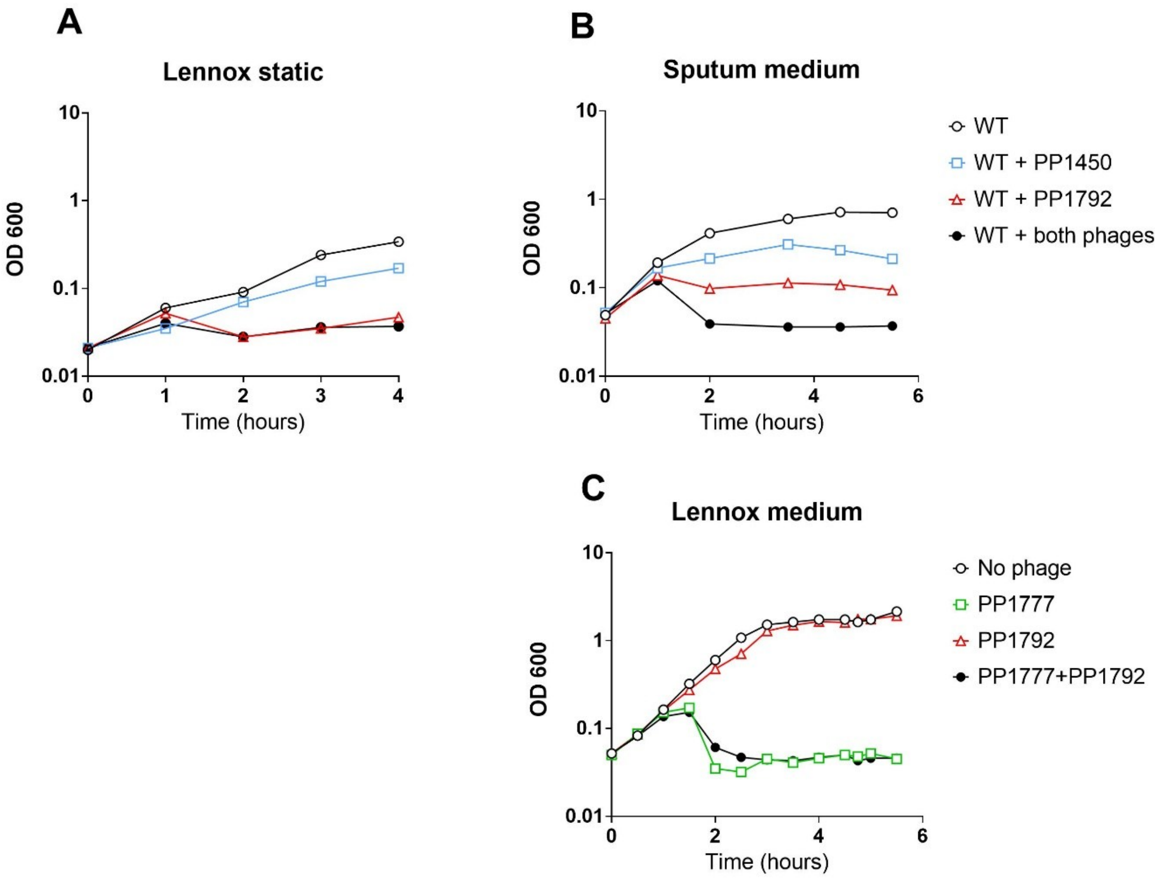

Figure S3: Immunoblot detection of Psl.

**A**: *Antibodies testing for the detection range*. Six serial 3-fold dilutions of the most concentrated Psl sample, which is a supernatant of the WT strain grown ON in Lennox. The top panel indicates the dilution factor of each sample deposited on the membrane. **B**: *Quantification of A,* dilutions below 0.04 are not detected, the background value corresponded to a volume of 30 000. All membranes for immunodetection contained this most concentrated sample as a reference, and images were recorded below the pixel saturation level of this sample. **C.** *Antibodies testing on negative controls*. Psl extractions were performed on strains MB119 (*pslD1* mutation) and MB132 (*pslA1* mutation) at the same OD tested for the WT strain, and samples were immunoblotted in parallel to the WT samples. Two independent extractions are shown. In addition, supernatants of the ON cultures of these two strains were also tested (one extraction shown). A signal close to the background was sometimes detected, but overall, the Psl antibodies were specific. **D**. *Psl detection with WT strain grown in MMMa*. Bound Psl from bacterial extractions (left, two independent extracts for each condition) and free Psl from supernatants (right, two independent extracts) are shown. **E**. *Psl detection with WT and MB79 strains grown in Lennox*. bound Psl from bacterial extractions (left, two independent extracts and blots) and free Psl from supernatants (right, two independent extracts and blots) are shown.

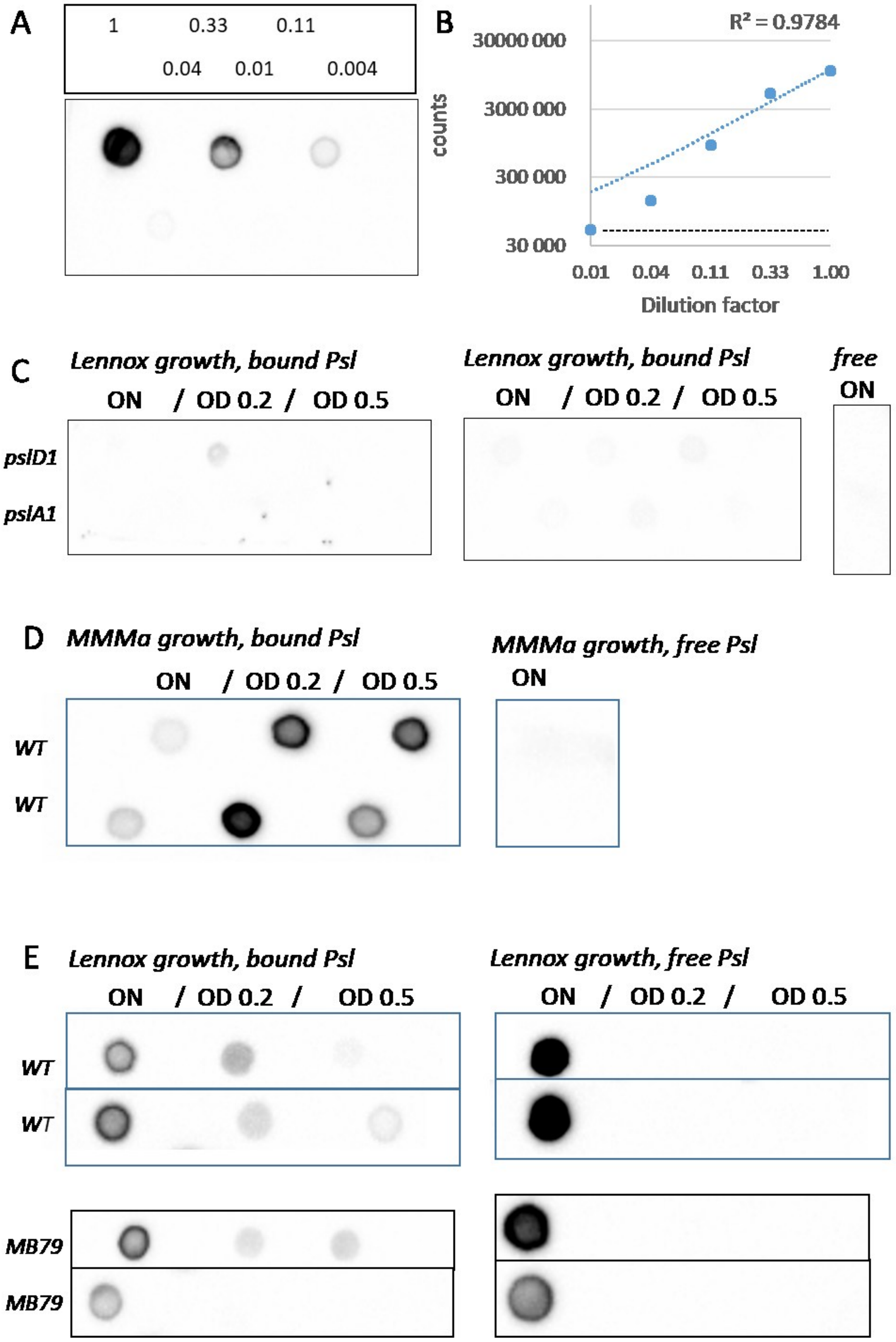
A
1
0.33
0.11
0.04
0.01
0.004
B
R² = 0.9784
30000 000
3000 000
300 000
30 000
counts
0.01
0.04
0.11
0.33
1.00
Dilution factor
C
Lennox growth, bound Psl
ON / OD 0.2 / OD 0.5
pslD1
pslA1
Lennox growth, bound Psl
ON / OD 0.2 / OD 0.5
free
ON
D
MMMa growth, bound Psl
ON / OD 0.2 / OD 0.5
WT
WT
MMMa growth, free Psl
ON
E
Lennox growth, bound Psl
ON / OD 0.2 / OD 0.5
WT
WT
Lennox growth, free Psl
ON / OD 0.2 / OD 0.5
MB79
MB79

Figure S4: **A**. Intubation device used for artificial respiration. **B**. Examples of crystal violet stained biofilms formed on a tubing piece of this device, after being incubated with PAO1-3Δ only (left), or successively with PAO1-3Δ (7 hours), then phages (16 hours).

A

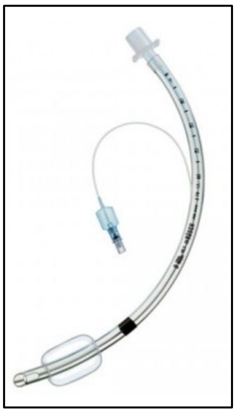

B

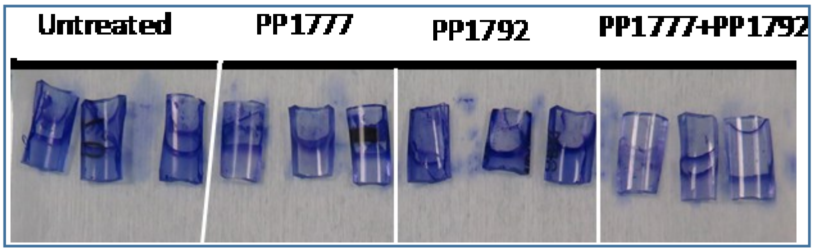

Figure S5: After 6 hours of PAO1_OR biofilm formation at 37°C in a 96-well plate, *LUZ24* phage PP1792 (5x10$^{6}$ PFU), *LS1* phage PP1450 (5x10$^{6}$ PFU), or a combination of both (10$^{7}$ in total) were added, and the biofilm growth or decay was monitored by confocal microscopy. Each image of the stack was then used to estimate the volume of viable bacteria at each time point, and volumes of living bacteria as a function of time are shown for three technical repetitions on the same plaque.

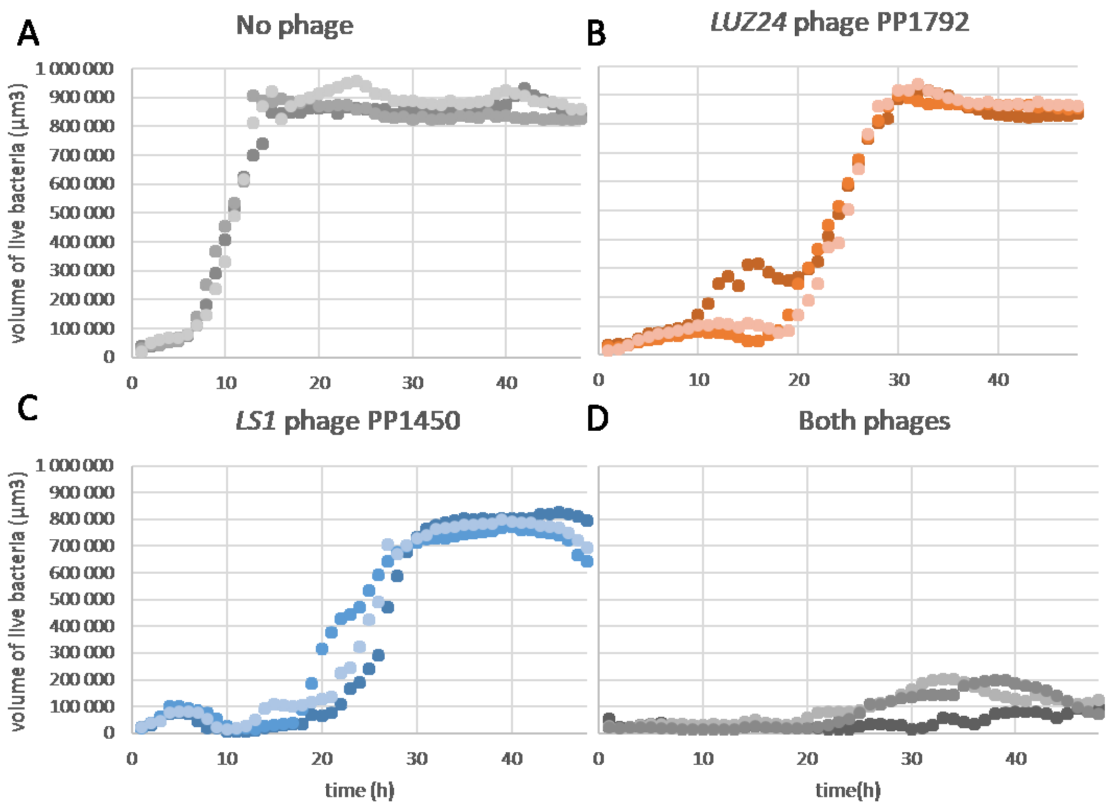

Figure S6: Second biological replicate of Figure 7.

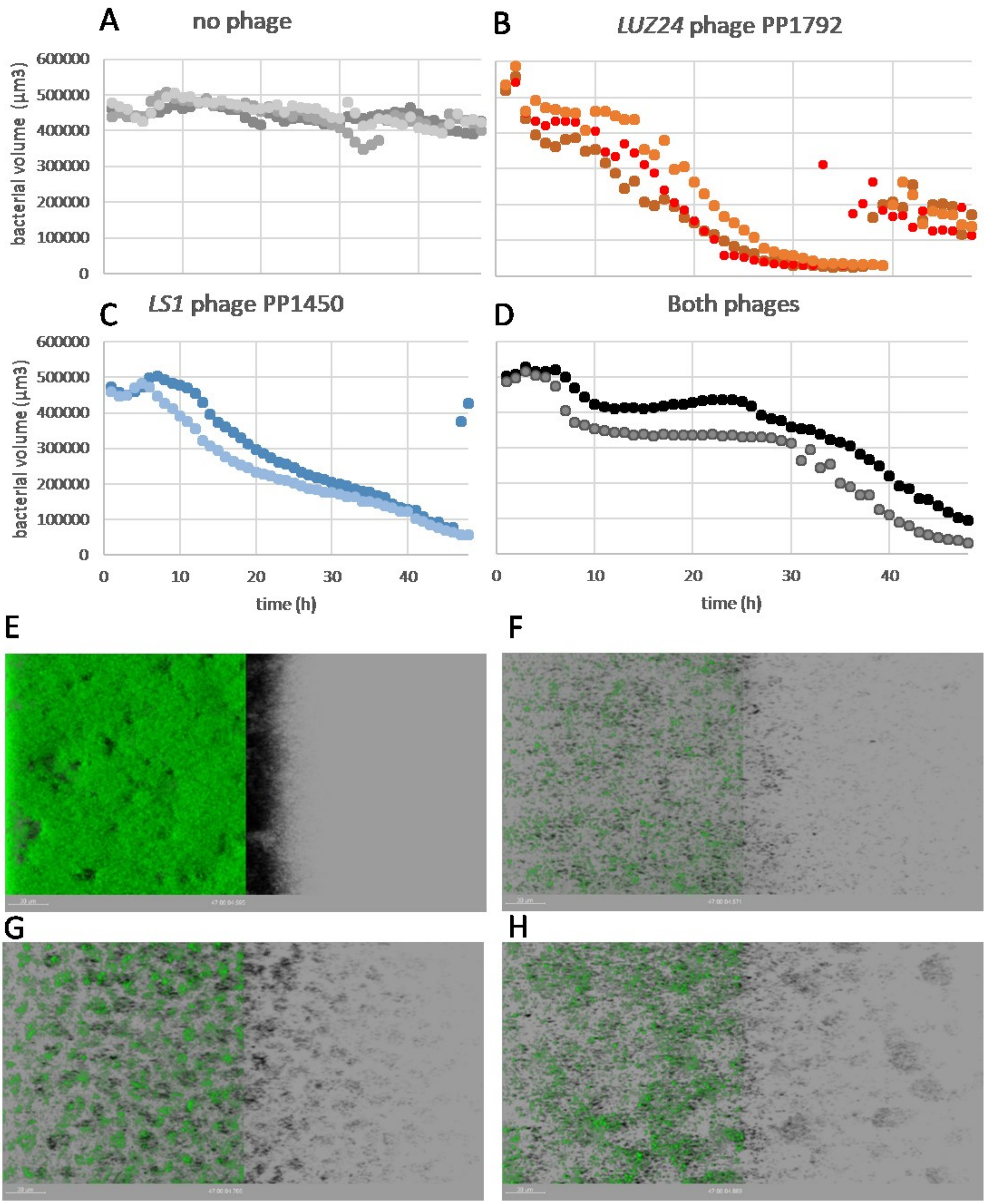

Figure S7: Third biological replicate of Figure 7.

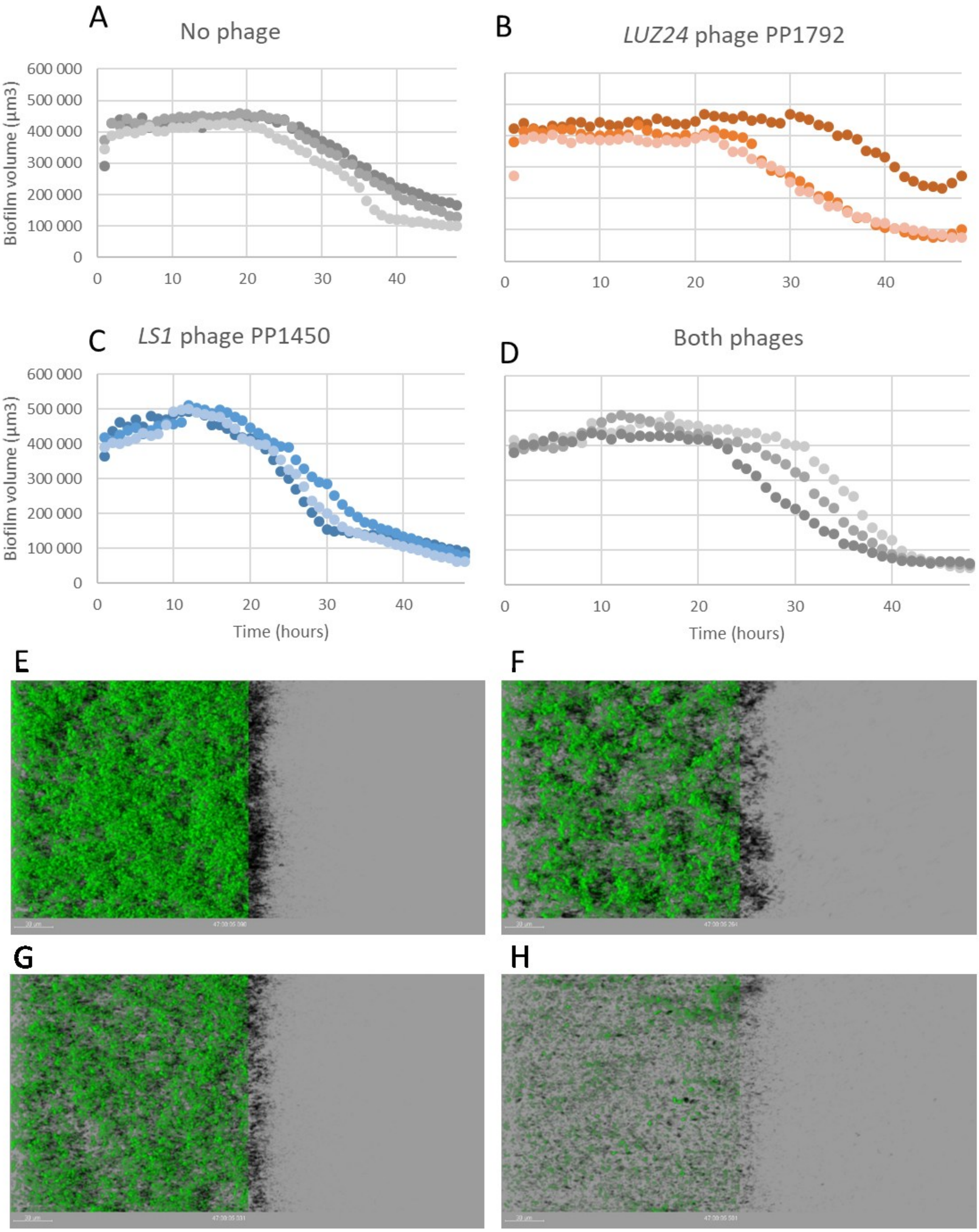

Figure S8: lytic activity of *LUZ24* phage PP1792 on the MB79 *wzy1* mutant strain, in Lennox medium. Growth conditions applied were as described in Figure 2. Phage added at MOI  1 at time 0.

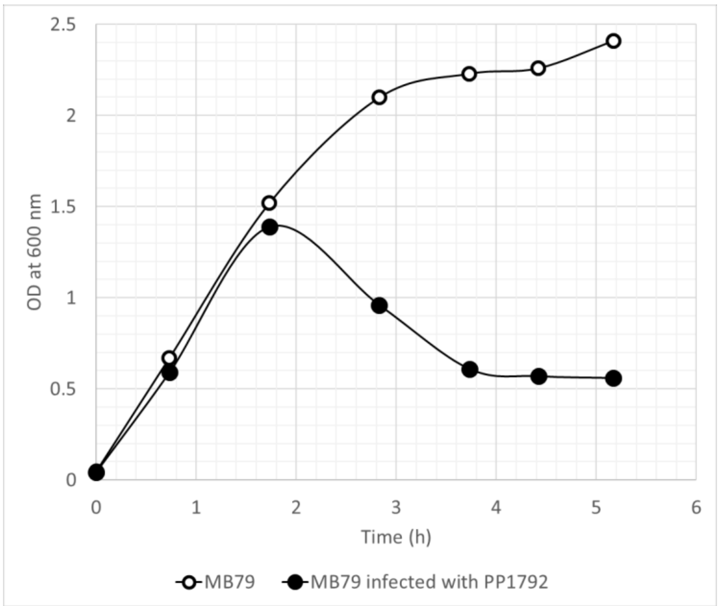

Figure S9: Psl amounts in the MB79 *wzy1* mutant strain, compared to the PAO1-3Δ strain (the PAO1-3Δ values are identical to Figure 5). Bacteria collected from ON cultures, or during exponential growth at OD 0.2 and 0.5 were used to extract surface-bound Psl from equivalent amounts of washed pellets (left side, "bound Psl"), and these extracts were then deposited on a membrane and treated for immunodetection as described in the Methods. In parallel, 5 μL of supernatants from the same cultures (right side, "free Psl") were directly deposited on the same membranes.

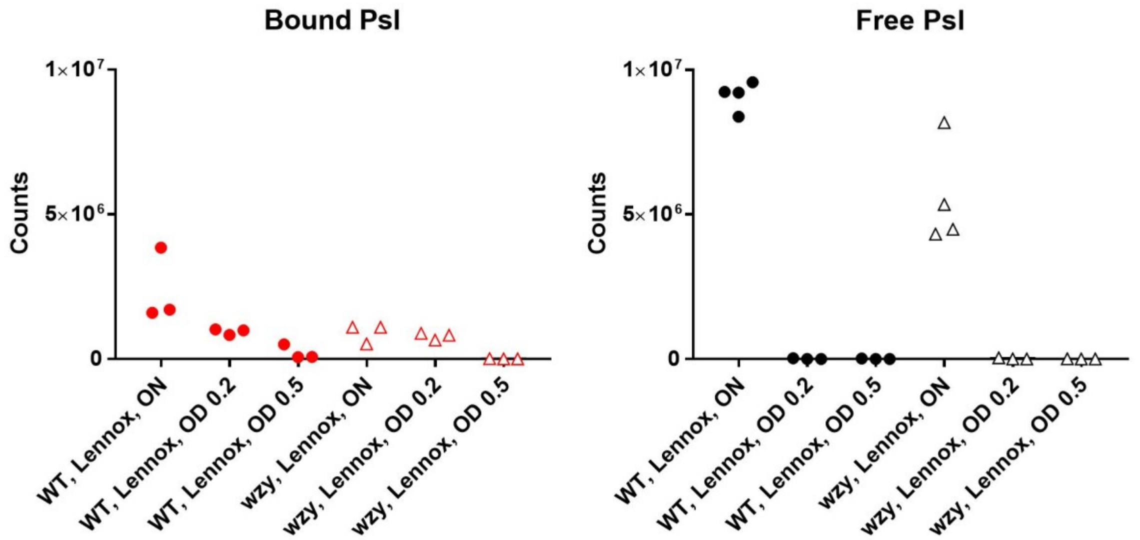

Figure S10. *LUZ24* phage PP1792 adsorption on *wzy1* derivatives in rich medium. Compared to the *wzy1* single mutant, double mutant strains MB87 and MB96 are not affected in their adsorption to phage PP1792 (One way ANOVA, P = 37 and 21%), whereas strain MB99 is an adsorption mutant (P< 0.01%). Adsorption on MB99 is also significantly diminished relative to PAO1-3Δ (P= 0.04%).

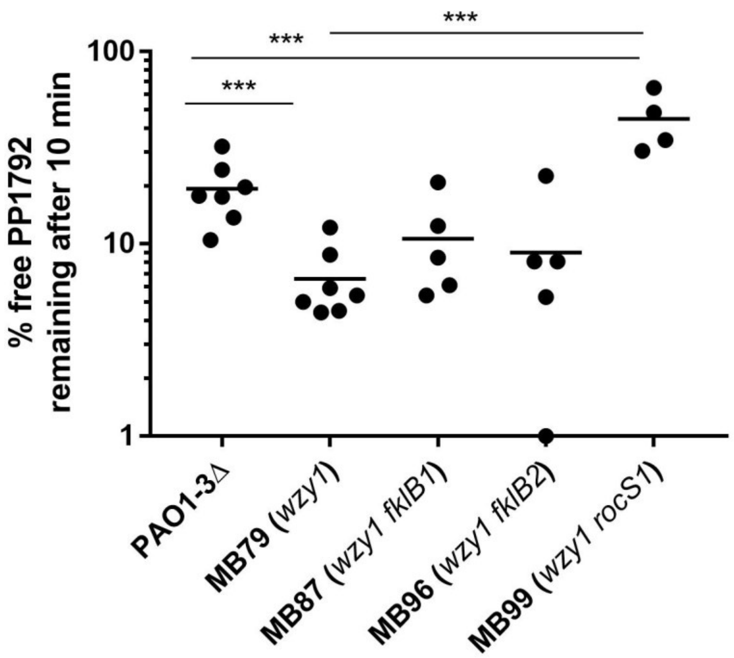
